# Large-scale mRNA translation and the intricate effects of competition for the finite pool of ribosomes

**DOI:** 10.1101/2021.09.15.460428

**Authors:** Aditi Jain, Michael Margaliot, Arvind Kumar Gupta

## Abstract

We present a new theoretical framework for large-scale mRNA translation using a network of models called the ribosome flow model with Langmuir kinetics (RFMLK), interconnected via a pool of free ribosomes. The input to each RFMLK depends on the pool density, and it affects the initiation rate and the internal ribosome entry rates at each site along each RFMLK. Ribosomes that detach from an RFMLK due to termination or premature drop-off are fed back into the pool. We prove that the network always converges to a steady-state, and study its sensitivity to variations in the parameters. For example, we show that if the drop-off rate at some site in some RFMLK is increased then the pool density increases and consequently the steady-state production rate in all the *other* RFMLKs increases. Surprisingly, we also show that modifying a parameter of a certain RFMLK can lead to arbitrary effects on the densities along the modified RFMLK, depending on the parameters in the entire network. We conclude that the competition for shared resources generates an indirect and intricate web of mutual effects between the mRNA molecules, that must be accounted for in any analysis of translation.

## 1 Introduction

Gene expression is a crucial cellular process that produces desired proteins by decoding the genetic code in the DNA [1]. In Eukaryotes, gene expression includes several processes. During transcription, the protein-coding information encoded in a DNA is copied into an mRNA molecule. During translation, the information inscribed in the mRNA is translated to chains of amino acids destined to become proteins [2,3]. The information encoded in the DNA and mRNA is decoded by complex molecular machines called RNA polymerases and ribosomes, respectively. The dynamical flow of these machines along the DNA or mRNA “tracks” plays an important role in gene expression and, in particular, in regulating protein production.

Translation is a fundamental process that takes place in all living cells, from bacteria to humans. Many mRNA molecules are translated in parallel with many ribosomes decoding every mRNA molecule [4, 5]. This implies that the mRNA molecules effectively “compete” for the finite resources in the cell, like tRNA molecules and free ribosomes [6, 7]. The competition for ribosomes may explain important dynamical properties of translation, that are difficult to understand when considering the translation of a single, isolated mRNA molecule. For example, it is known that stalling ribosomes may detach from the mRNA before completing the translation process [8, 9]. This is somewhat surprising, as a ribosome that drops-off from the mRNA before reaching the stop codon fails to complete the synthesis of a full-length protein, and releases a truncated protein, whose accumulation could be detrimental to the cell [10]. However, in the context of competition for free ribosomes, premature drop-off may have a positive effect: it allows stalled ribosomes to join the pool of free ribosomes that can initiate translation in other mRNA molecules. Thus, modeling translation as a network of interconnected processes and taking into account competition for shared resources is important for gaining a deeper understanding of fundamental principles in cellular biophysics.

Typically, translation is initiated by ribosome scanning from the 5′ end of the capped mRNA. However, some mRNAs include internal ribosome entry sites (IRESs), that allow for translation initiation in a cap-independent manner. IRESs, first discovered in poliovirus, are common in RNA viruses and allow viral translation even when host translation is inhibited for some reason [11, 14]. Cellular growth regulatory genes, and genes transcribed in response to stress also contain IRES elements [12]. Also, synthetic biologists often insert IRES sequences into their vectors to allow expressing two or more genes from a single vector [15]. We believe that the effect of IRESs on translation should also take into account the competition for shared resources, like ribosomes. For example, a recent study [13] shows that the non-structural protein 1 (Nsp1), produced by the SARS-CoV-2 virus, binds to the human 40S subunit in ribosomal complexes, and thus interferes with mRNA binding. Since translation of the viral mRNA is more efficient than that of cellular mRNAs with 5′ UTRs, the net effect is hijacking the cellular translation machinery by the virus.

Many mathematical and computational models for translation on a single mRNA molecule have been developed [16]. One such model is the Asymmetric Simple Exclusion Process (ASEP), and its variants motivated by biological findings [17–23]. This model includes a 1D chain of sites, and particles that hop according to a stochastic mechanism from a site to a neighboring site, provided the latter site is empty. Thus, every site is either free or includes a single particle. This simple exclusion principle represents the constraint that two particles cannot be in the same place at the same time. If motion is unidirectional, then ASEP is called the totally asymmetric exclusion process (TASEP). In the context of translation, the lattice represents the mRNA molecule and the hopping particles are the ribosomes. ASEP has been used extensively to model gene translation, and various other multi-agent systems with local interactions including intracellular transport, pedestrian dynamics, and more [24, 25].

Despite its simple description, analysis of ASEP is non-trivial because the simple exclusion principle generates an intricate coupling between the hopping particles. In particular, analysis results for TASEP are asymptotic, i.e. they hold when the number of sites converges to infinity, and closed-form results exist only under the restrictive assumption that all the internal hopping rates are equal [26].

A deterministic mathematical model called the Ribosome Flow Model (RFM), which can be derived as a dynamical mean-field approximation of TASEP, was suggested in [27]. This model is highly amenable to analysis using tools from systems and control theory. The RFM, and its various generalizations, have been used to model and analyze sophisticated features of translation, for example, ribosome flow along a circular mRNA [29], the effect of ribosome recycling [30], a ribosome flow model with different site sizes [32], ribosome flow incorporating bidirectional flow and the phenomena of attachment and drop-off [35], entrainment of the protein production rate to periodic elongation rates [36], and regulating the average protein synthesis in the cell under stochastic variability in the ribosome elongation rates [37]. Networks of interconnected RFMs have also been studied [38].

Raveh et al. [33] analyzed a model called the RFM network with a pool (RFMNP), which includes a network of RFMs, interconnected via a pool of free ribosomes. For a recent application of this model to ribosome traffic engineering, see [34]. However, the RFMNP cannot model the important features of premature drop-off and IRESs. As we will see below, adding these features to the model generates new, important and perhaps surprising results.

In this paper, we consider a network of RFMs with an additional Langmuir kinetics called RFMLKs. This allows modeling drop-off and attachment of ribosomes at any site along the mRNA due to premature drop-off or IRESs, respectively. The RFMLKs are interconnected via a pool of free ribosomes yielding a new model referred to as the *RFM with Langmuir kinetics network* (RFMLKN). This allows modeling simultaneous translation of an arbitrary number of mRNA molecules, with ribosome drop-off from any site along the mRNA molecule to the pool of free ribosomes, and attachment at an IRES at any site along the mRNA. In particular, we use this model to rigorously analyze the effect of ribosome drop-off and/or IRES in one mRNA on the production rate of all the other mRNA molecules.

We use the powerful theory of strictly cooperative dynamical systems with a first integral [39] to prove that the RFMLKN admits a continuum of linearly ordered equilibrium points. Every solution of the system converges to an equilibrium that depends on the network parameters and the total number of ribosomes in the network. This represents a dynamical steady-state where the ribosome flow into and out of every site along any mRNA molecule is equal, and the flows into and out of the pool are also equal. Thus, any two solutions starting from two initial conditions corresponding to an equal total number of ribosomes in the network converge to the same equilibrium point. In other words, the network “forgets” the exact initial condition, except for the total initial density of ribosomes. This qualitative behaviour holds for any feasible set of parameters covering many possible biophysical conditions.

We also show that if all the transition rates vary in a periodic manner, with a common period *T*, then every solution of the RFMLKN converges to a periodic solution with period *T*. This implies in particular that the protein production rate entrains to periodic excitations in the translational machinery.

We use the RFMLKN to analyze quantitatively and qualitatively important questions such as the effect of ribosome drop-off/attachment from/to a site in one mRNA on the steady-state production rate of all the other mRNAs in the network. The analysis highlights how the competition for shared resources generates an indirect and intricate web of mutual effects between the mRNA molecules in the cell. These effects cannot be analyzed using models of translation on a single, isolated mRNA molecule.

The remainder of this paper is organized as follows. The next section summarizes the main analysis results and their biological implications. Section 3 presents the new mathematical model and demonstrates using several examples how it can be used to study large-scale translation in the cell. Section 4 states our main theoretical results. For the sake of readability, all the proofs are placed in the Appendix. The final section concludes and suggests possible directions for further research.

## 2 Summary of Main Results and their Biological Implications

The RFMLKN encapsulates many fundamental aspects of gene translation. During mRNA translation, ribosomes attach at the 5′ end of the mRNA and scan it in a sequential manner. At each elongation step, every sequence of three consecutive nucleotides in the mRNA, called a codon, is decoded into an amino-acid, and this process continues until the ribosome reaches the 3′ end of the mRNA [2]. The codon decoding rates may vary among different mRNAs and depend on many transcript features [40]. Several ribosomes may scan the same mRNA molecule in parallel, but a ribosome cannot overtake another ribosome in front of it, thus obeying the simple exclusion principle. Ribosomes may detach from the mRNA molecule before reaching the stop codon due to several reasons like ribosome “traffic jams”, the presence of a premature stop codon, ribosome-ribosome interactions due to depletion of aminoacyl tRNA or amino-acid misincorporation, etc. [41]. The limited availability of free ribosomes induces indirect coupling due to competition between mRNA molecules.

We prove that for a given set of elongation, drop-off and attachment rates, and a total number of ribosomes in the network, the RFMLKN admits a unique steady-state i.e. the ribosomal density profiles on all the mRNAs and in the pool converge to a fixed value, as time goes to infinity. This raises the important question of how does the steady-state changes if we modify any parameter in the model, e.g. the rate of ribosome drop-off in one site of a specific mRNA.

Our analysis shows that following an increase (decrease) in a drop-off rate in any mRNA molecule, the steady-state ribosome density profile in all the *other* mRNA molecules increases (decreases). The intuitive explanation for this is as follows: increasing the drop-off rate leads to releasing more ribosomes to the pool of free ribosomes and this increases the initiation rate as well as the attachment rate in all the other mRNA molecules leading to an increase in their ribosome density profile. We also prove the “dual” result, namely, that increasing (decreasing) an attachment rate in a specific mRNA decreases (increases) the steady-state ribosome densities in all the *other* mRNA molecules. However, and perhaps surprisingly, we show that it is very difficult to predict the effect of a variation in one of the rates on the mRNA that is modified, as the effect will depend on the entire network. For example, increasing the attachment rate in one site of a specific mRNA may deplete the pool and thus decrease the effective attachment rates in other sites along the modified mRNA, leading to an unexpected decrease in the density along this mRNA.

These results highlight the indirect effects of competition for resources, and also the importance of taking competition into account when “tinkering” with the bio-physical features of a single mRNA molecule e.g. by replacing codons by synonymous codons.

Our simulations suggest another interesting implication of ribosome drop-off and/or ribosome attachment (e.g. in IRESs). It seems that these phenomena increase the amount of indirect “communication” between the mRNA molecules, through the pool, and thus lead to a higher level of synchronization between the production rates in the mRNAs. This suggests another possible advantage of ribosome drop-off and attachment as tools for regulating the protein production from different copies of the same mRNA.

## 3 Mathematical Model

Our model is a network of interconnected ribosome flow models with Langmuir kinetics (RFMLK). We begin by reviewing the RFMLK and then describe the network of interconnected RFMLKs.

### 3.1 Ribosome Flow Model with Langmuir Kinetics (RFMLK)

The RFMLK is a deterministic, non-linear, continuous-time compartmental model for modeling ribosome flow along a single mRNA molecule and is a coarse-grained mean field approximation of TASEP with Langmuir kinetics and open boundary conditions [28, 35].

The RFMLK describes the flow along *n* consecutive sites. In the context of translation, each site corresponds to a codon or a group of codons along the mRNA molecule. The flow from site *i* to site *i* + 1 is determined by a positive parameter *λ*_*i*_, with units 1/time. The flow from site *i* to the environment [from the environment to site *i*] is determined by a non-negative parameter *α*_*i*_ [*β*_*i*_], with units 1/time. The state variable *x*_*i*_(*t*) : ℝ_+_ → [0, 1], *i* = 1, 2, … *n*, describes the normalized density of ribosomes at site *i* at time *t*. This may also be interpreted as the probability that site *i* is occupied at time *t*. The dynamical equations describing the RFMLK are:

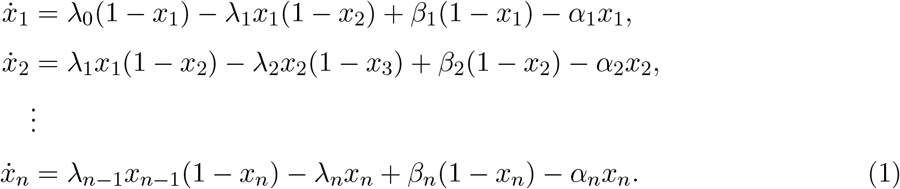

Defining *x*_0_(*t*) ≡ 1 and *x*_*n*+1_(*t*) ≡ 0 allows to write these equations in the more succinct form:

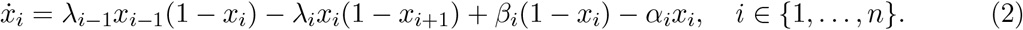

The term *λ*_*i−*1_*x*_*i*−1_(1 − *x*_*i*_) represents the flow of the particles from site *i* − 1 to site *i*. This increases with the density at site *i* − 1 and decreases as site *i* becomes fuller. This corresponds to a “soft” version of the simple exclusion principle in ASEP. Similarly, the term *λ*_*i*_*x*_*i*_(1 − *x*_*i*+1_) represents the flow from site *i* to *i* + 1. The term *β*_*i*_(1 − *x*_*i*_) represents the attachment of particles from the environment to site *i*, whereas *α*_*i*_*x*_*i*_ represents the drop-off of particles from site *i* to the environment. By setting some of the *α*_*i*_s and *β*_*i*_s to positive values and the others to zero, it is possible to model drop-off and attachment at specific sites. If *α*_*i*_ = *β*_*i*_ = 0 for all *i* then the model reduces to the RFM. The topology of the RFMLK is depicted in Figure 1.

**Figure 1:**
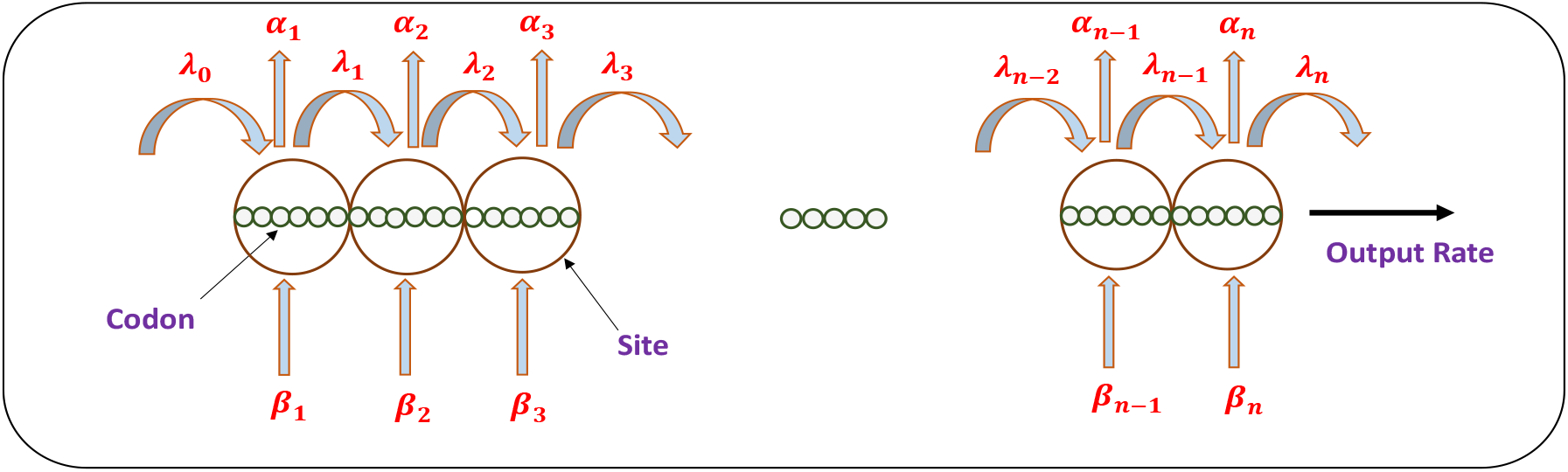
The RFMLK models unidirectional flow along a chain of *n* sites. The density at site *i* at time *t* is represented by *x*_*i*_(*t*). The parameter *λ*_*i*_ > 0 controls the transition rate from site *i* to site *i* + 1, with *λ*_0_ and *λ*_*n*_ controlling the initiation and termination rates, respectively. The parameter *α*_*i*_ ≥ 0 [*β*_*i*_ ≥ 0] controls the drop-off [attachment] rate from [to] site *i*.

**Figure 2:**
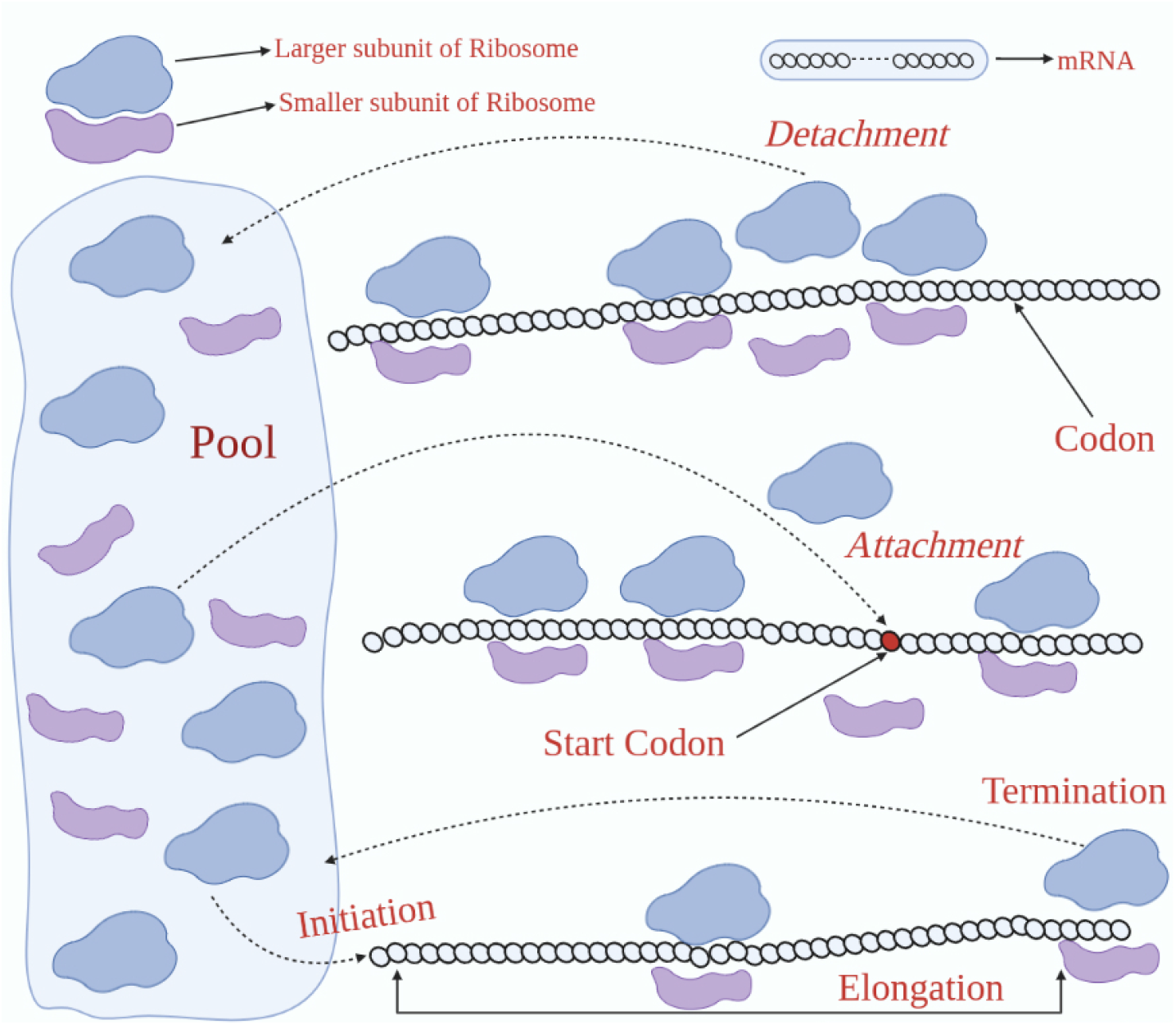
Large-scale translation of mRNA molecules in the cell. Several ribosomes may decode the same mRNA. Ribosomes that detach from an mRNA enter the pool of free ribosomes.

To build a network, we use an RFMLK with an input and output given by

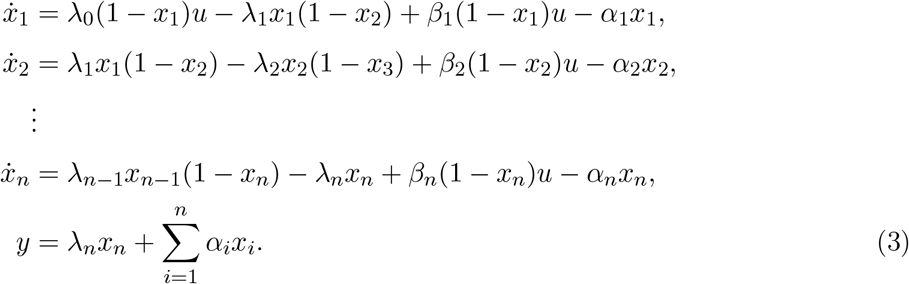

The time-varying control *u*(*t*) multiplies the term representing the entry rate into site 1, and also the attachment rates in all the sites. We assume that *u*(*t*) ≥ 0 for any time *t*. A larger value of *u*(*t*) corresponds for example to a larger density of free ribosomes in the vicinity of the mRNA at time *t*, and consequently it increases the effective initiation rate in the first site and the attachment rates in all the sites. The output *y*(*t*) is the total exit rate of ribosomes from the RFMLK to the environment at time *t*.

Note that (3) is a nonlinear model, as it includes both products of state-variables and products of state-variables and the control input.

#### Example 1

Consider the case *n* = 1. In this special case, (3) becomes the linear system

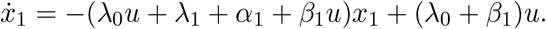

Fix *x*(0) ∈ [0, 1]. Consider a constant control *u*(*t*) ≡ *v*, with *v* > 0, then it is clear that *x*(*t*) ∈ (0, 1) for all *t* > 0, and that the limit *e*_1_ := lim_*t*→∞_ *x*_1_(*t*) exists, and satisfies

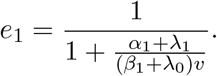

In particular, if *v* = 0 then *e*_1_ = 0, and if *v* → ∞ then *e*_1_ → 1. The first case corresponds to no ribosomes in the vicinity of the mRNA, so the single site empties. The second case corresponds to an infinite density of ribosomes, so the site fills up completely. Note also that *e*_1_ is an increasing function of *λ*_0_, *β*_1_, *v*, and a decreasing function of *λ*_1_, *α*_1_.

The next subsection introduces the RFMLKN.

### 3.2 A network of Ribosome Flow Models with Langmuir Kinetics and a pool

To model competition for the finite pool of ribosomes in the cell, we consider a set of *m* RFMLKs with input and output, representing *m* different mRNA molecules in the cell, connected via a pool of free ribosomes.

The *i*th RFMLK has length *n*_*i*_, input function *u*^*i*^, output *y*^*i*^, and rates 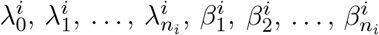 and 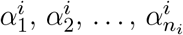. The dynamics of the *m* RFMLKs is written as

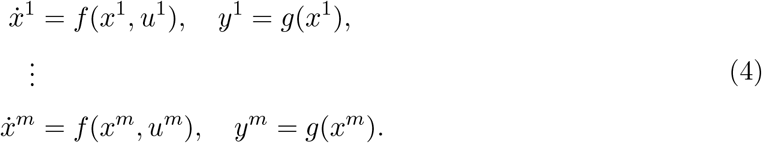

These RFMLKs are interconnected through a pool of free ribosomes, i.e. ribosomes that are not attached to any mRNA. We use the scalar function *z*(*t*) ≥ 0 to denote the density of ribosomes in the pool at time *t*. The pool feeds the initiation locations as well as the sites in the mRNAs where attachment takes place. Mathematically, this implies that *u*^*i*^(*t*) = *G*_*i*_(*z*(*t*)), *i* = 1, 2, …, *m*. We assume that every function *G*_*i*_(·) : ℝ_+_ → ℝ_+_ satisfies the following properties:

1. *G*_*i*_(0) = 0;
2. *G*_*i*_(·) is continuously differentiable with 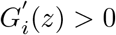 for all *z* ≥ 0; and
3. There exists *c* > 0 such that *G*_*i*_(*z*) ≤ *cz* for all *z* > 0 sufficiently small.

The first property implies that if the pool is empty then no ribosomes can exit the pool; the second implies that as the number of ribosomes in the pool increases, more ribosomes exit the pool; and the third is a technical condition that is needed for the proof given later on of persistence in the RFMLKN. Functions that satisfy these properties include, for example, the linear function *G*(*z*) = *az*, with *a* > 0, and the bounded function *G*(*z*) = *a* tanh(*bz*), with *a, b* > 0 [20, 42].

The dynamics of the *i*th RFMLK in the network is thus given by:

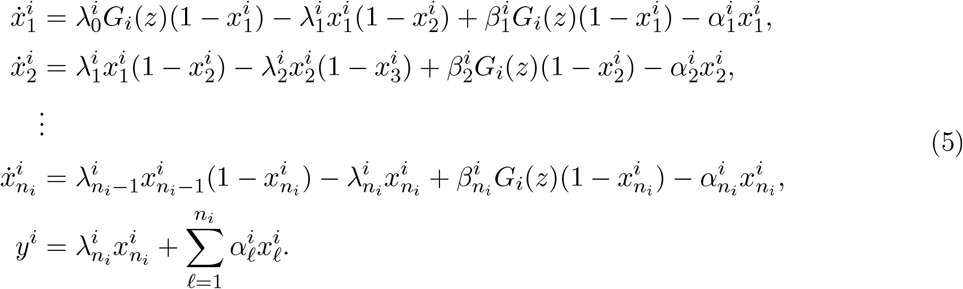

The output of each RFMLK is fed into the pool. Hence, the pool dynamics is given by:

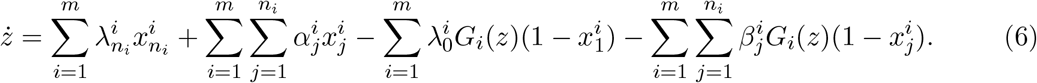

In other words, all the ribosomes that exit or drop-off the mRNAs feed the pool, and the pool feeds the initiation and attachment sites in all the mRNAs.

Summarizing, the RFMLKN is a dynamical system with 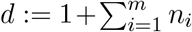 state variables, and dynamics described by equations (4), (5) and (6) (see Figure 3). Eq. (6) and our assumptions on the functions *G*_*i*_ imply that if *z*(0) ≥ 0 then *z*(*t*) ≥ 0 for all *t* ≥ 0, i.e. the pool density is always non-negative.

**Figure 3:**
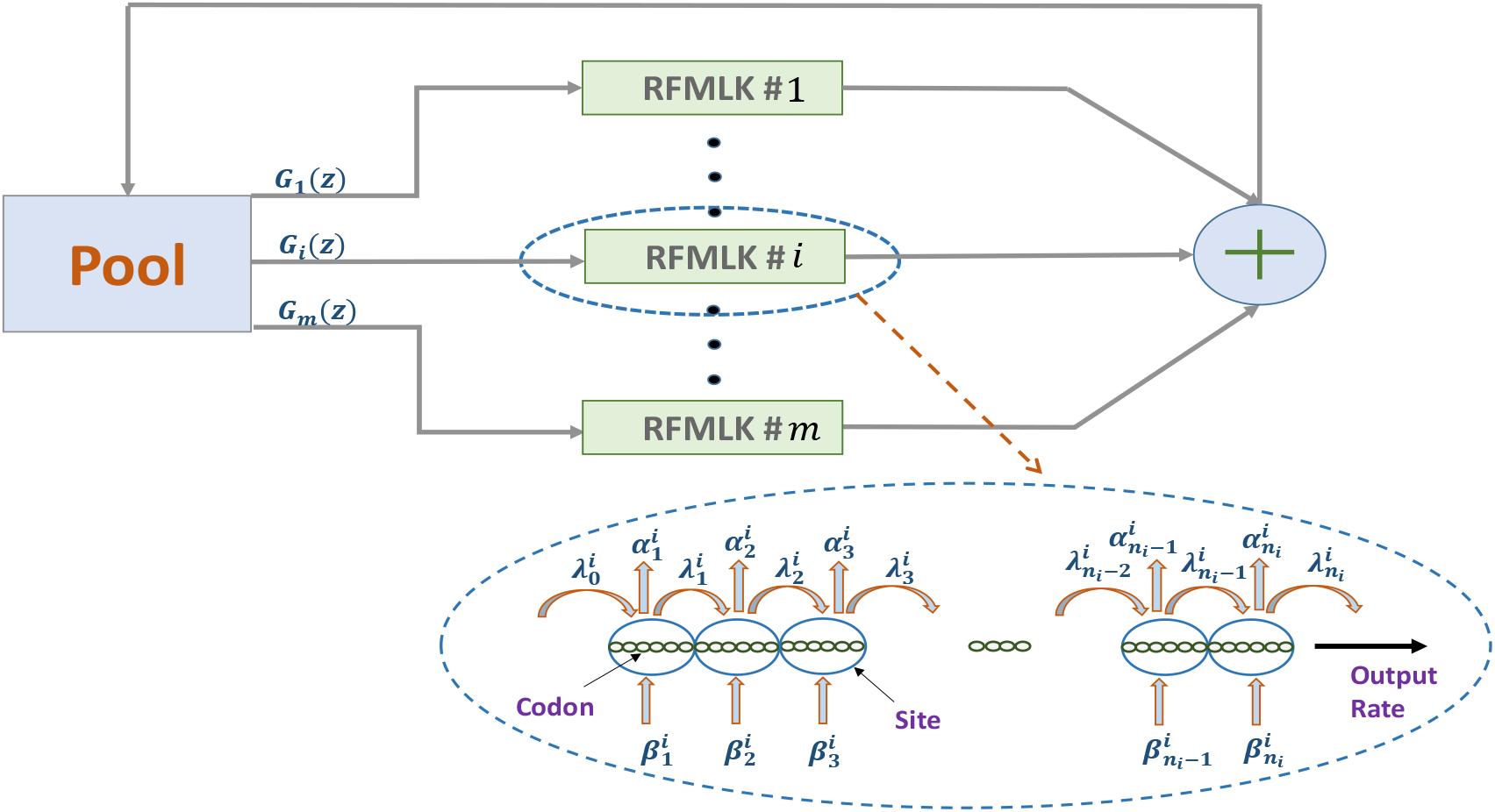
Each mRNA is described by an RFMLK with input and output. The output of each RFMLK is fed into the pool, and the pool feeds the initiation and attachment rates in all the RFMLKs.

Let

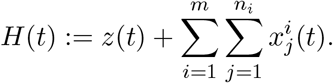

This is the total number of ribosomes in the system at time *t*. An important property of the RFMLKN is that it is a closed system, so *H*(*t*) is conserved, i.e. *H*(*t*) ≡ *H*(0) for all *t* ≥ 0. Thus, *H* is a first integral of the dynamics.

The RFMLKN models the indirect coupling between the mRNA molecules induced by competition for the finite number of ribosomes in the system. For example, if there is a “traffic jam” of ribosomes on one of the mRNAs then the pool is depleted and thus the initiation and attachment rates to all the mRNAs decrease.

We prove in Section 4 that all the state variables in the RFMLKN converge to a steady-state. The steady-state values depend on the parameter values in all the RFMLKs and the total number of ribosomes in the network. Let 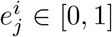 denote the steady-state density at the *j*th site in the *i*th RFMLK, and let *e*_*z*_ ∈ [0, ∞) denote the steady-state pool density.

The RFMLKN provides a versatile and powerful framework for simulating and analyzing the effect of various biological phenomena on steady-state translation in the cell under competition for free ribosomes. In the examples below, we demonstrate how various changes in the network affect the RFMLKN steady-state. We also explain the biological implications of the simulation results.

Our first example demonstrates how the total number of ribosomes in the system affects the ribosomal densities in the mRNAs.

#### Example 2

Consider an RFMLKN that includes a single RFMLK with dimension *n*_1_ = 3, rates 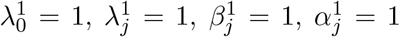, for *j* = 1, 2, 3, and a pool with an output function *G*(*z*) = tanh(*z*). We simulated this system with the initial condition 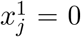 for all *j*, and *z*(0) = *c*, so that the total number of ribosomes in the system is *c*, for various values of *c*. When *c* = 0 there are no ribosomes in the network and the steady-state values are all zero. As *c* increases, the number of ribosomes along the RFMLK increases. Since tanh(*z*) → 1 as *z* → ∞, the initiation and attachment rates converge to 1. Thus, ribosomal densities in RFMLK saturate to the values corresponding to initiation rate 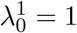, and attachment rates 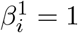. The remaining ribosomes accumulate in the pool (see Figure 4a). Using a different output function, namely, *G*(*z*) = *z*, we see in Figure 4b that as *c* increases, all ribosomal densities tend to the maximal possible value 1, that is, the RFMLK completely fills up and the remaining ribosomes accumulate in the pool.

**Figure 4:**
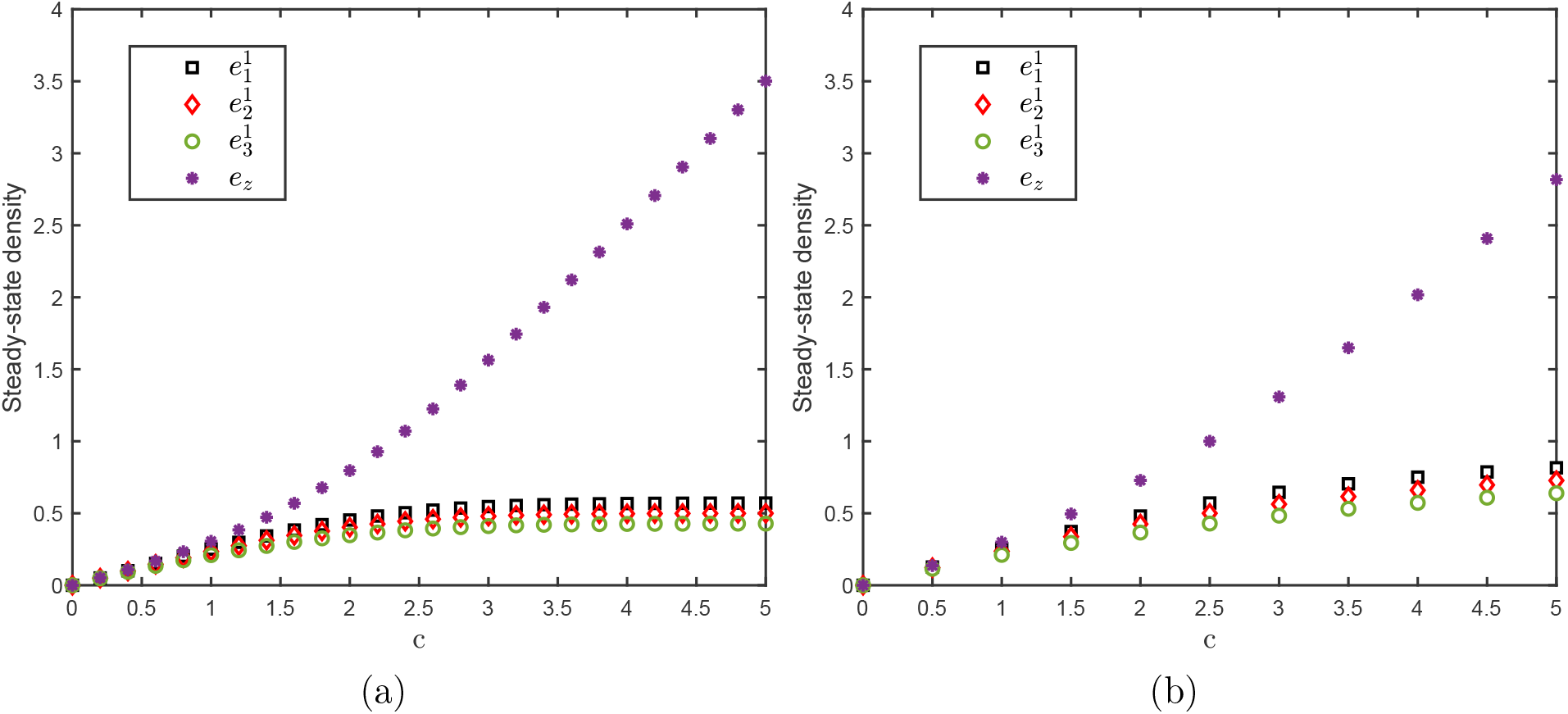
a) Steady-state values in the RFMLKN in Example 2 as a function of the total number of ribosomes *c*. (a) when *G*(*z*) = tanh(*z*); (b) when *G*(*z*) = *z*.

The next example describes the effect of the drop-off rate of ribosomes along a coding region in one of the mRNA molecules on the steady-state profiles of all the mRNAs in the network. Let

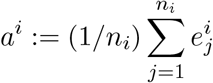

denote the average steady-state density (ASSD) in the *i*th RFMLK.

#### Example 3

Consider an RFMLKN with *m* = 3 RFMLKs with dimensions *n*_*i*_ = 3, *i* = 1, 2, 3, and parameters 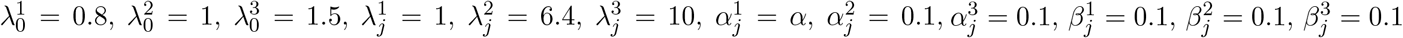, for all *j*, and *G*_*i*_(*z*) = *z, i* = 1, 2, 3. The initial condition is 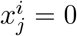, for all *i, j*, and *z*(0) = 2. We vary the parameter *α*, i.e. the ribosome drop-off rate from all the sites in the first RFMLK, in the range [0.1, 3]. Figure 5 depicts the ASSD in each RFMLK as a function of *α*. It can be seen that as *α* increases, the ASSD in the first RFMLK decreases, whereas the ASSD in all the other RFMLKs increases. Indeed, as the drop-off rate from the first RFMLK increases, the density in the pool increases, and more ribosomes become available for translating the other mRNA molecules, thus increasing the ASSD in the other RFMLKs.

**Figure 5:**
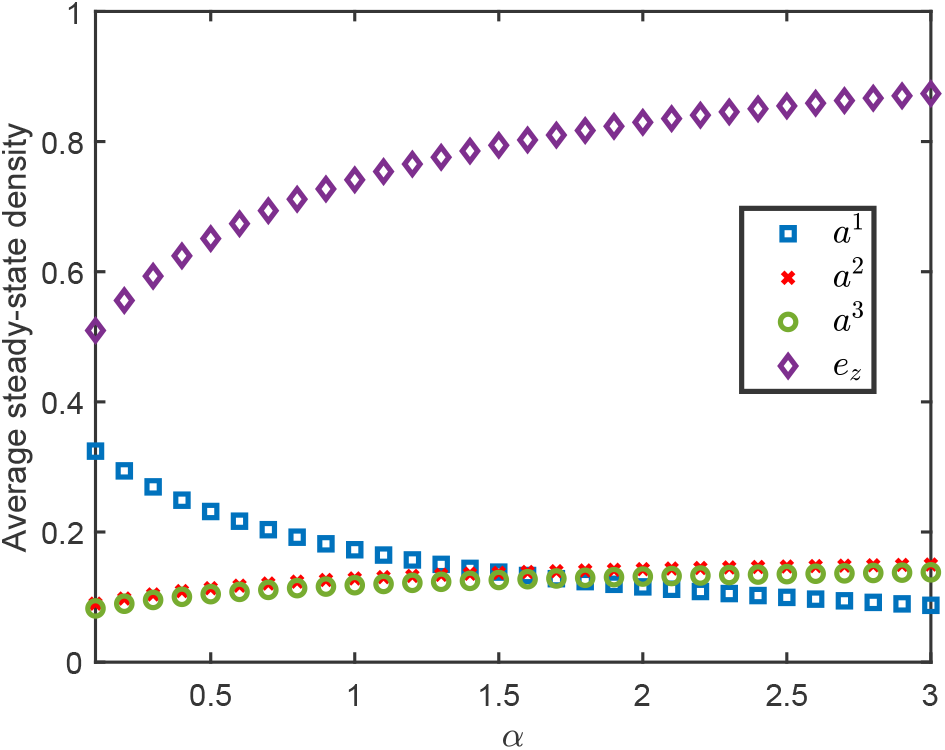
Average steady-state density in the RFMLKN in Example 3 as a function of the drop-off rate *α* in the first RFMLK.

From a biological perspective, this example corresponds to a situation when due to genetic errors or insufficient availability of charged tRNAs or frameshifting [43], ribosomes start detaching before reaching the stop codon in an mRNA, resulting in truncated protein products. Our results explain why this may still be beneficial to the cell. The ribosome drop-off from one mRNA molecule increases the number of free ribosomes that are now available to translate other mRNAs which in turn increases the corresponding protein production rates.

The next example considers the “dual” case of increasing the attachment rate in one of the mRNA molecules in the network.

#### Example 4

Consider an RFMLKN with *m* = 2 RFMLKs with dimensions *n*_*i*_ = 10, *i* = 1, 2. The parameter values are 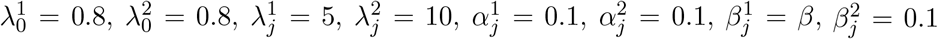, for all *j*, and *G*_*i*_(*z*) = tanh(*z*), *i* = 1, 2. The initial condition is 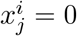, for all *i, j*, and *z*(0) = 3.5. Figure 6 depicts the ASSD in each RFMLK as a function of *β* ranging from 0.1 to 3. It can be seen that as *β* increases, the ASSD in the first RFMLK increases and the ASSD in the other RFMLK decreases. This is due to the attachment of ribosomes at the first RFMLK leading to a depletion of ribosomes in the pool, and thus to a decrease in the ASSD in the second RFMLK. Note the relatively sharp decrease in the steady-state pool density as *β* increases. This is due to the fact that the number of sites is *n*_*i*_ = 10, so a “traffic jam” in an RFMLK involves many stalled ribosomes along the RFMLK.

**Figure 6:**
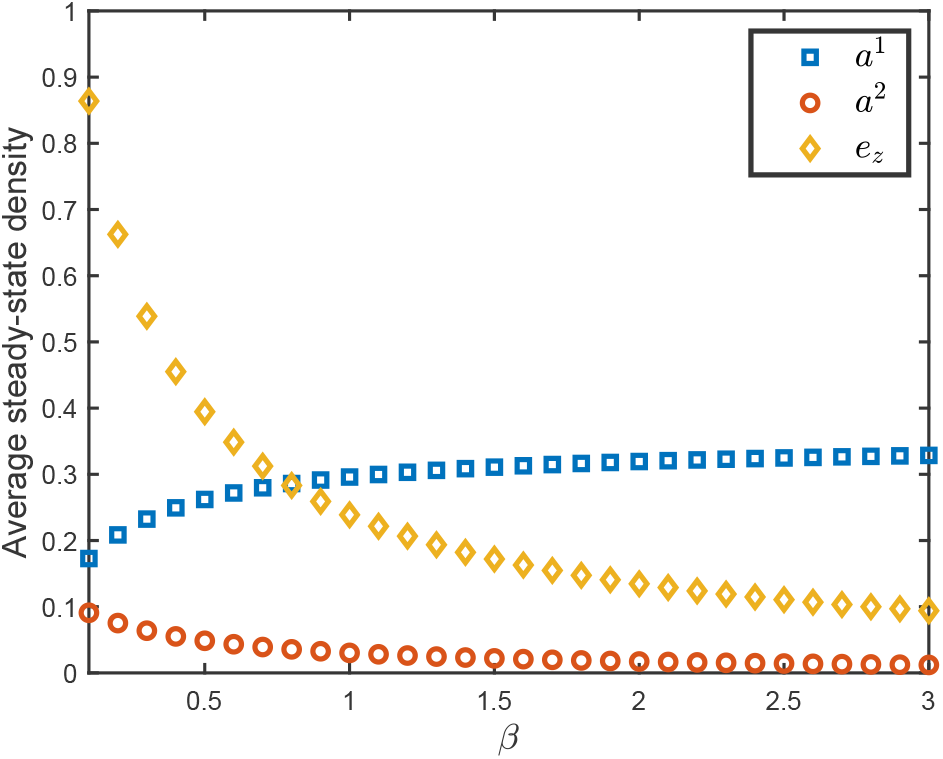
Average steady-state density in the RFMLKN in Example 4 as a function of the attachment rate *β* in the first RFMLK.

From a biological point of view, the attachment rate models internal ribosome entry sites (IRESs) that are common for example in viral mRNA. The RFMLKN may thus be used to shed more light on how the viral mRNA “overtakes” the pool of free ribosomes and thus: (1) accelerates the translation of viral mRNA, and (2) concomitantly slows down the cellular innate immune response [47]. IRESs have also been used as a biotechnological tool allowing the synthesis of several proteins of interest from one multicistronic mRNA [44–46]. In this context, the example above shows that the design of such tools should also take into consideration their effect on the pool of free ribosomes.

The next example demonstrates the effect of modifying the length of one mRNA molecule in the network.

#### Example 5

Consider an RFMLKN with *m* = 2 RFMLKs with dimensions *n*_1_ = 5, and *n*_2_, respectively. The parameter values are 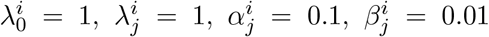, for all *i, j*, and *G*_*i*_(*z*) = tanh(*z*), *i* = 1, 2. The initial condition is 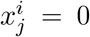, for all *i, j*, and *z*(0) = 5. We simulated this network for various values of *n*_2_. As *n*_2_ increases, there is a decrease in the ASSD in both RFMLKs and in the pool density (see Figure 7). Indeed, increasing *n*_2_ implies that ribosomes that bind to the second RFMLK remain on it for a longer period of time. This decreases the steady-state pool density and, consequently, the steady-state densities in all the RFMLKs.

**Figure 7:**
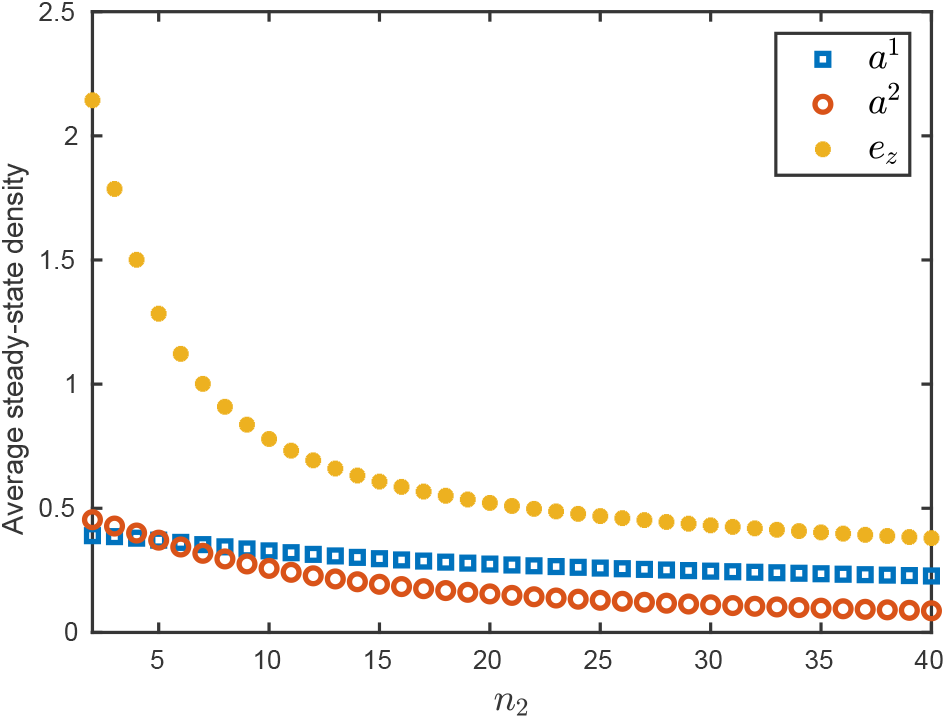
ASSD and *e*_*z*_ in Example 5 as a function of the length *n*_2_ of the second RFMLK.

The next example again studies the effect of increasing *n*_2_ and also compares the RFMLKN and the RFMNP.

#### Example 6

Consider an RFMLKN with *m* = 2 RFMLKs with dimensions *n*_1_ = 5, and *n*_2_, respectively. The parameters are 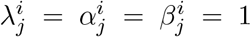 for all *i, j*, and the initial condition is 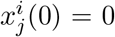 for all *i, j* and *z*(0) = 25. Recall that 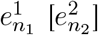 denotes the steady-state density in the last site of the first [second] RFMLK. Let 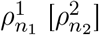 denote the steady-state density in the last site of the first [second] RFMLK, when 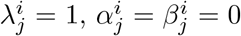 for all *i, j*, so the RFMLKs reduce to RFMs. Figure 8 shows that as *n*_2_ increases, the steady-state densities 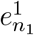 and 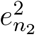 tend to equal values. However, 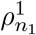 and 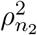 are different. The reason for this may be that the non-zero attachment and detachment rates increase the indirect communication between the RFMLKs (through the pool) leading to better “synchronization”.

**Figure 8:**
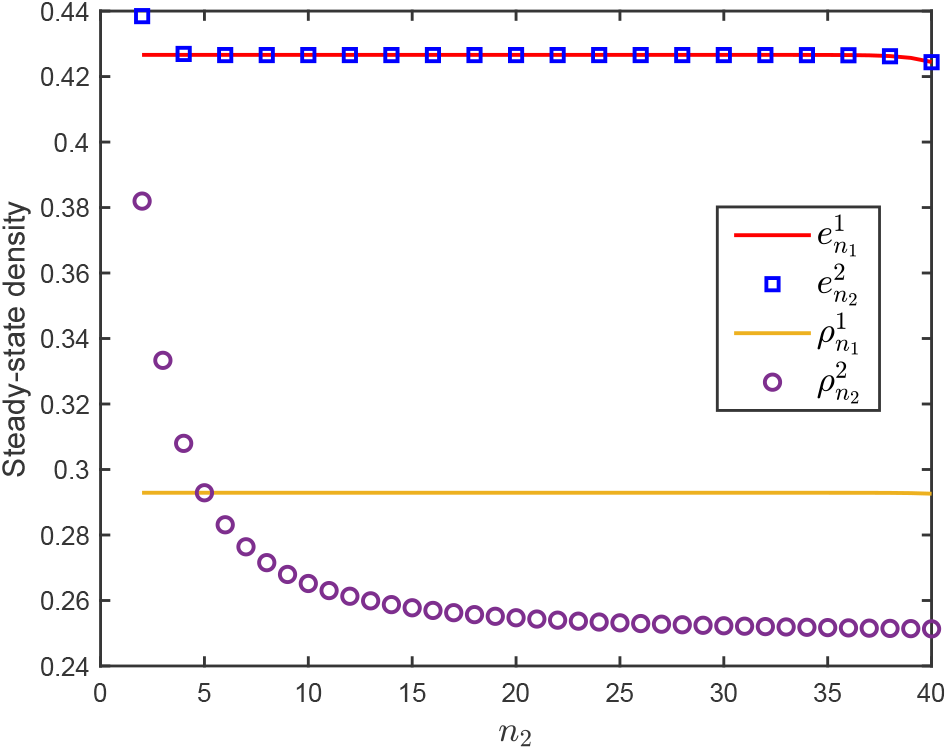
Steady-state densities in Example 6 as a function of the length *n*_2_ of the second RFMLK.

The next section rigorously analyzes the RFMLK and the RFMLKN using tools from systems and control theory and in particular the theory of cooperative dynamical systems [48].

## 4 Main Results

We begin by analyzing the properties of the RFMLK with input and output described in (3), as these are the basic ingredients of the RFMLKN. Recall that *x*_*i*_(*t*) ∈ [0, 1] for all *t*, so the state-space of the RFMLK is *C*^*n*^ := [0, 1]^*n*^. Let int(*C*^*n*^) and *∂C*^*n*^ denote the interior and boundary of *C*^*n*^ respectively. Let *x*(*t, a*) denote the solution of equation (3) at time *t* ≥ 0 for the initial condition *a* ∈ *C*^*n*^. For the sake of readability, all the proofs are placed in the Appendix.

### 4.1 Persistence

If *u*(*t*) ≡ 0 then no ribosomes enter the RFMLK and then it is clear that *x*(*t*) will converge to zero, that is, the density of ribosomes at each site will go to zero. The next result shows that for any input that is bounded below by a positive number all the state-variables remain bounded away from zero and also bounded away from one.

#### Proposition 4.1

*Consider the RFMLK with a control u such that u*(*t*) ≥ *s* > 0 *for all t* ≥ 0. *For any τ* > 0 *there exists ϵ* = *ϵ*(*τ*) > 0, *with ϵ*(*τ*) → 0 *as τ* → 0, *such that for any initial condition a* ∈ *C*^*n*^ *the solution of* (3) *satisfies*

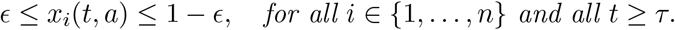

In other words, for any control *u*(*t*) that is strictly positive for all *t* ≥ 0 we have that after any time *τ* > 0 all the normalized densities are strictly separated away from zero and from one. In particular, if the densities converge to a steady-state *e* then *e*_*i*_ ∈ (0, 1) for all *i*.

### 4.2 Contraction

Contraction theory is a powerful tool for analyzing nonlinear dynamical systems [57, 58], and has found applications in bio-molecular systems, control theory, synchronization of coupled non-linear systems [59], reaction-diffusion differential equations [60], and more.

For *x* ∈ ℝ^*n*^, let |*x*|_1_ := |*x*_1_| + · · · + |*x*_*n*_| denote the *L*_1_ norm of *x*. For a non-singular matrix *P* ∈ ℝ^*n×n*^, let |*x*|_*P*,1_ := |*Px*|_1_, i.e. the scaled *L*_1_ norm of *x*.

#### Proposition 4.2

*Consider the RFMLK with a control u such that u*(*t*) ≥ *s* > 0 *for all t* ≥ 0, *and fix τ* > 0. *There exist a non-singular matrix P* = *P* (*τ*) *and η* = *η*(*τ*) > 0 *such that for any a, b* ∈ *C*^*n*^, *we have*

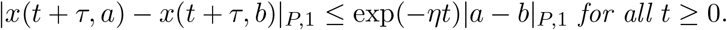

In other words, the RFMLK is contracting with respect to the scaled norm | · |_*P*,1_ after (the arbitrarily small) time delay *τ*. Proposition 4.2 implies several useful asymptotic properties of the RFMLK. These are described in the following subsections.

### 4.3 Global asymptotic stability

#### Proposition 4.3

*The RFMLK with a constant control u*(*t*) ≡ *s* > 0 *admits a unique steady-state e*^*s*^ ∈ int(*C*^*n*^) *that is globally asymptotically stable, i*.*e*.,

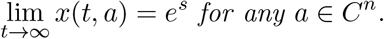

In other words, the solution converges to *e*^*s*^ for any initial condition, so the inital condition is “forgotten”. The equilibrium *e*^*s*^ represents a dynamic steady-state where the input and output flows from each site in the RFMLK are equal, and thus the densities in each site are constant.

### 4.4 Monotone control system

Angeli and Sontag [31] extended the notion of a monotone system to control systems. The next result shows that the RFMLK is a monotone control system. For two vectors *v, w* ∈ ℝ^*n*^, we write *v* ≤ *w* if *v*_*i*_ ≤ *w*_*i*_ for all *i* = 1, …, *n*, and *v* ≪ *w* if *v*_*i*_ *< w*_*i*_ for all *i* = 1, …, *n*.

#### Proposition 4.4

*Fix two initial conditions a, b* ∈ *C*^*n*^, *with a* ≤ *b, and two controls u, v* : ℝ_+_ → ℝ_+_, *with u*(*t*) ≤ *v*(*t*) *for all t* ≥ 0. *Then the corresponding solutions of the RFMLK satisfy*

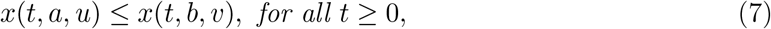

*and*

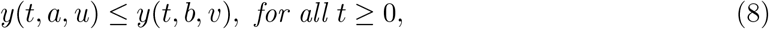

In other words, if we consider two identical RFMLKs–the first with initial densities *a*_*i*_ and the second with initial densities *b*_*i*_, with *a*_*i*_ ≤ *b*_*i*_ for all *i*, and apply a control *u* in the first and *v* in the second, with *u*(*t*) ≤ *v*(*t*) for all *t* ≥ 0–then at each time *t* ≥ 0 each density in the first RFMLK will be smaller or equal to the corresponding density in the second RFMLK.

The next proposition analyzes the relation between the steady-state densities corresponding to constant control values.

#### Proposition 4.5

*Consider the RFMLK with constant controls u*(*t*) ≡ *s*^1^ *and v*(*t*) ≡ *s*^2^ *with* 0 *< s*^1^ *< s*^2^. *Fix a, b* ∈ *C*^*n*^, *and let*

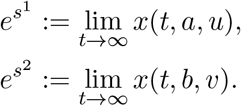

*Then*

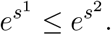

We now turn to analyze the RFMLKN. For a matrix 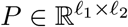, let 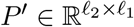 denote the transpose of *P*. Recall that every 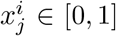, and that the pool density satisfies *z* ∈ [0, ∞), so the state-space of the RFMLKN is

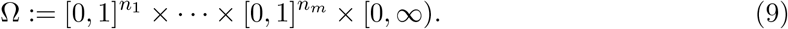

For *a* ∈ Ω, let 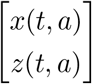 denote the solution of the RFMLKN at time *t* with the initial condition *a*. Let *d* := *n*_1_ + · · · + *n*_*m*_ + 1, and let 1_*d*_ denote a column vector of *d* ones. For *s* ≥ 0, let 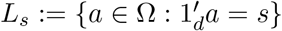, i.e. the *s* level set of the first integral *H*. In other words, *L*_*s*_ is the set of all states in Ω with a total density of ribosomes equal to *s*.

### 4.5 Invariance and persistence

The next result follows immediately from the equations of the RFMLKN.

#### Proposition 4.6

*The state space* Ω *in* (9) *is an invariant set for the dynamics of the RFMLKN that is, if a* ∈ Ω *then* 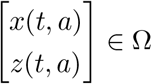 *for all t* ≥ 0.

In other words, every trajectory emanating from an initial condition in the state space remains in it for all *t* ≥ 0.

The next result shows that trajectories that begin from an initial condition in Ω become uniformly separated from the boundary of Ω.

#### Proposition 4.7

*Consider the RFMLKN. For any τ* > 0 *there exists ϵ* = *ϵ*(*τ*) > 0, *with ϵ*(*τ*) → 0 *as τ* → 0, *such that for any a* ∈ Ω \ {0} *we have*

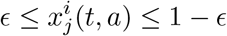

*and*

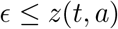

*for all t* ≥ *τ, i* ∈ {1, 2, …, *m*}, *and j* ∈ {1, 2 …, *n*_*i*_}.

In other words, after an arbitrarily short time every density in every RFMLK is in the range [*ϵ*, 1 − *ϵ*], and the pool density is in [*ϵ*, ∞). To explain why this result is useful, note that the Jacobian *J* of the vector field of the RFMLK with input and output satisfies *J* (*x, u*) = *M* (*x*) − *D*(*x, u*), where *D*(*x, u*) is a diagonal matrix with entries

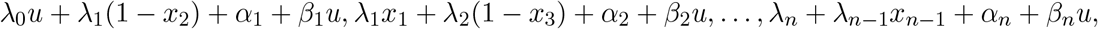

and

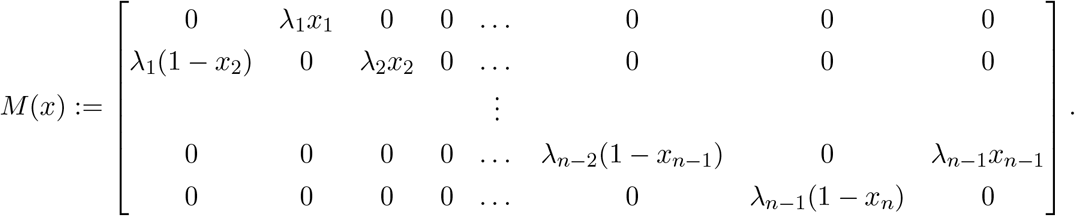

For any *x* ∈ [0, 1]^*n*^ all the entries of *M* (*x*) are nonnegative, so (5) is a cooperative dynamical system [54]. The matrix *M* (*x*) may become reducible for values *x* on the boundary of [0, 1]^*n*^. However, *M* (*x*) is irreducible for all *x* ∈ (0, 1)^*n*^. Thus, Proposition 4.7 guarantees that after an arbitrarily short time the matrix *M* (*x*(*t*)) and, thus *J* (*x*(*t*), *u*(*t*)), becomes an irreducible matrix. This will be used in analyzing the asymptotic properties of the RFMLKN described below.

### 4.6 Stability

The next result shows that every level set contains a unique steady-state ribosome distribution in each mRNA and in the pool. The proof is based on the theory of monotone dynamical systems that admit a first integral, see [55, 56].

#### Theorem 4.1

*Every level set L*_*s*_ *contains a unique equilibrium point* 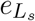 *of the RFMLKN and for any initial condition a* ∈ *L*_*s*_, *the solution of the RFMLKN converges to* 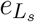. *Furthermore, for any* 0 ≤ *s < p*,

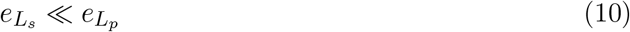

In other words, the RFMLKN admits a continuum of equilibrium points and any two solutions starting from initial conditions in the same level set of the system converge to the same equilibrium point. Thus, the rates 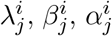 and the total number of ribosomes *s* in the network determine a unique steady-state density profile in the RFMLKs and the pool. Equation (10) implies that for two initial conditions, the first one with a smaller total number of ribosomes than the second one, the corresponding equilibrium points *e*^1^ and *e*^2^ will be completely ordered: every steady-state density in *e*^1^ will be strictly smaller than the corresponding density in *e*^2^.

#### Example 7

To model a gene that is highly expressed with respect to other genes, consider an RFMLKN with a single RFMLK and a pool. Assume that the output function of the pool is *G*_1_(*z*) = *z*, and that the dimension of the RFMLK is *n*_1_ = 2. The equations of the RFMLKN are then

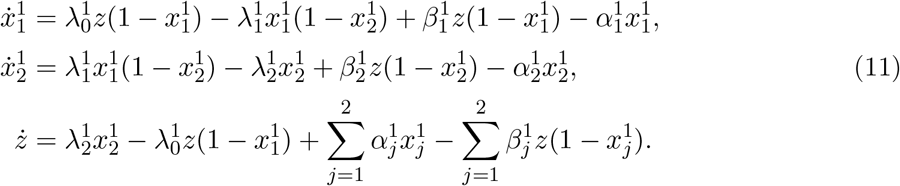

Any equilibrium point 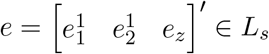 satisfies

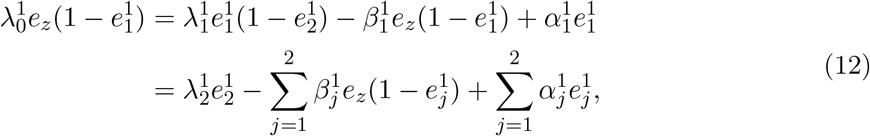

and

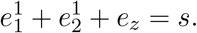

Figure 9 depicts trajectories of (11) with parameters 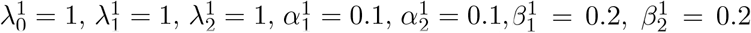, and three different initial conditions in the level set *L*_1_: [0.5 0.5 0]′, [0.5 0 0.5]′, and [0 0.5 0.5]′. It may be seen that the three solutions converge to the same equilibrium point.

**Figure 9:**
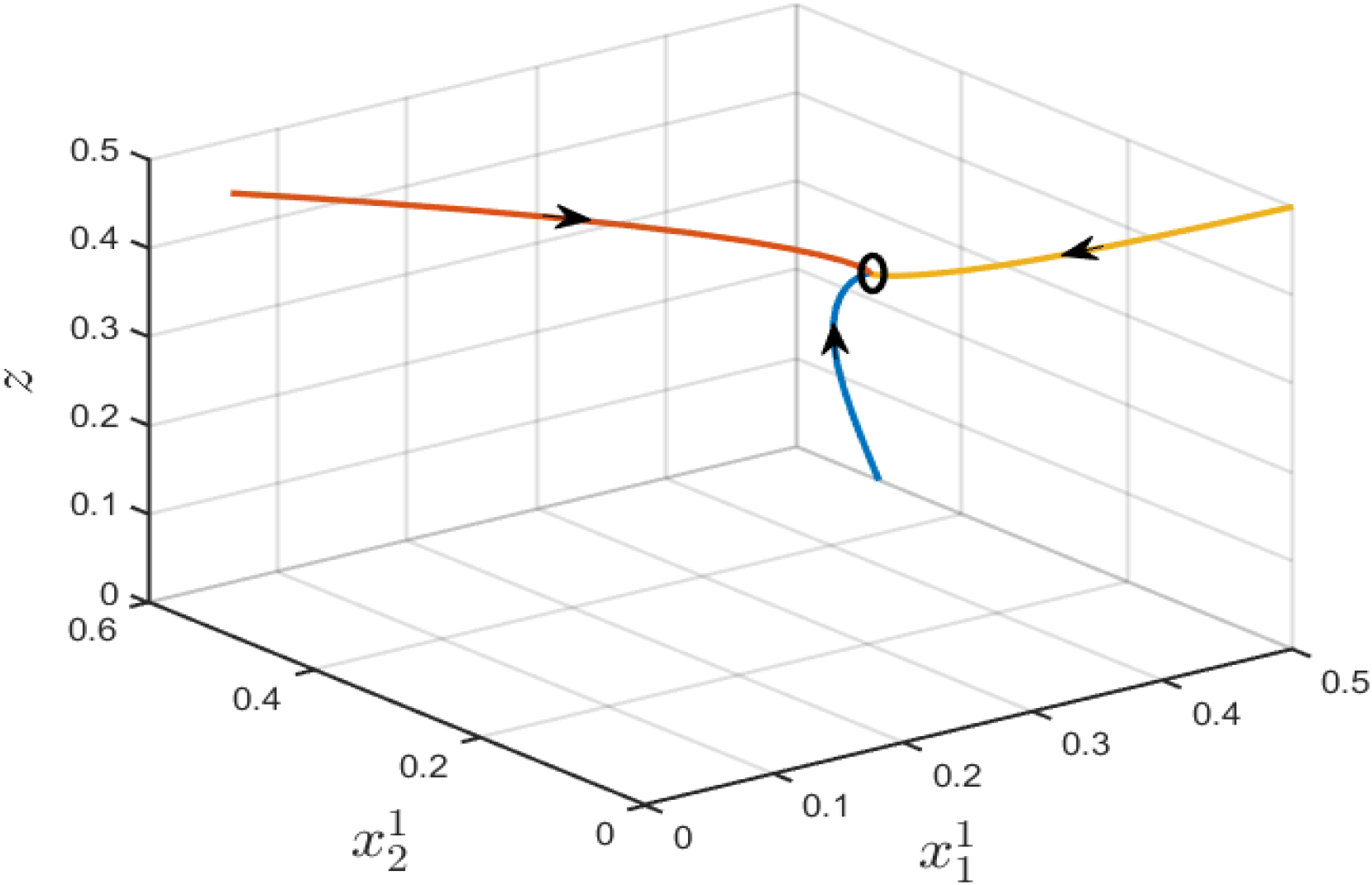
Trajectories of the RFMLKN in Example 7 for three initial conditions in *L*_1_. The unique equilibrium in *L*_1_ is marked by a circle.

Various intracellular mechanisms may affect the parameters of the translation machinery. For example, the elongation rates depend on the interaction between the nascent peptide and the exit tunnel of the system [49]. Stress conditions increase ribosome abortion and drop-off. Due to competition for the finite pool of ribosomes, any change in the translation speed along a specific mRNA molecule will also indirectly affect the translation of other mRNAs in the network. The variability in the factors that affect translation in the cell requires models that can be used to analyze the sensitivity to parameter values. Another motivation for studying these issues comes from synthetic biology, for example, the recent interest in co-expression of multiple genes at a given, desired ratio [50, 51].

### 4.7 Effect of parameters

Our first result in this subsection analyzes the affect of a modification in the drop-off rate at one site of an mRNA molecule on the entire RFMLKN. We assume, without loss of generality, that the modification is in one of the rates in the first RFMLK.

#### Theorem 4.2

*Consider an RFMLKN with m RFMLKs with dimensions n*_*i*_, *i* = 1, …, *m, and parameters* 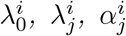, *and* 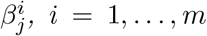 *and j* = 1, 2, …, *n*_*i*_. *Fix s* > 0, *and let* 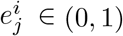, *and e*_*z*_ ∈ (0, ∞) *denote the unique steady-state density in the level set L*_*s*_ *of H. Fix k* ∈ {1, …, *n*_1_}, *and suppose that we modify the RFMLKN by changing* 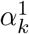 *to*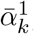, *with* 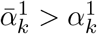. *Let* 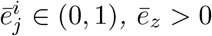, *denote the steady-state density in the modified RFMLKN. Then*

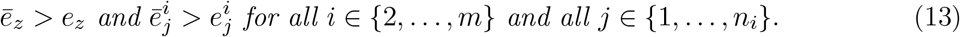

In other words, an increase in the drop-off rate in one of the RFMLKs yields an increase in the steady-state pool density and consequently an increase in the density in each site in all the *other* RFMLKs.

The effect of increasing 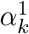 on the steady-state in the first RFMLK is non-trivial. It is natural to expect a decrease in the density in each site of the first RFMLK. But, as more ribosomes accumulate in the pool the effective attachment rates in sites along the first RFMLK may also increase, leading to an increase in the density in certain sites. In general, the total effect on the first RFMLK will depend on all the parameters in the RFMLKN, and is thus difficult to predict. The next two examples demonstrate this.

#### Example 8

Consider an RFMLKN with *m* = 2 RFMLKs of dimensions *n*_1_ = 6 and *n*_2_ = 3, parameters 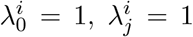, for all 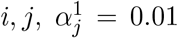, for *j* = 1, 2, 4, 5, 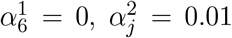, for *j* = 1, 2, 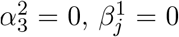, for all 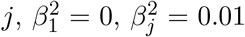, for *j* = 2, 3, and *G*_*i*_(*z*) = *z, i* = 1, 2. The initial condition is 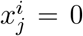, for all *i, j*, and *z*(0) = 5. We simulated this RFMLKN for several values of the drop-off rate 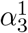 from the third site in RFMLK #1. Figures 10a and 10b show that increasing 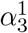 increases 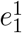, and decreases 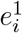 for all *i* > 1. As predicted in Thm. 4.2, it also increases *e*_*z*_, so more ribosomes accumulate in the pool leading to an increase in 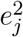 for all *j*.

**Figure 10:**
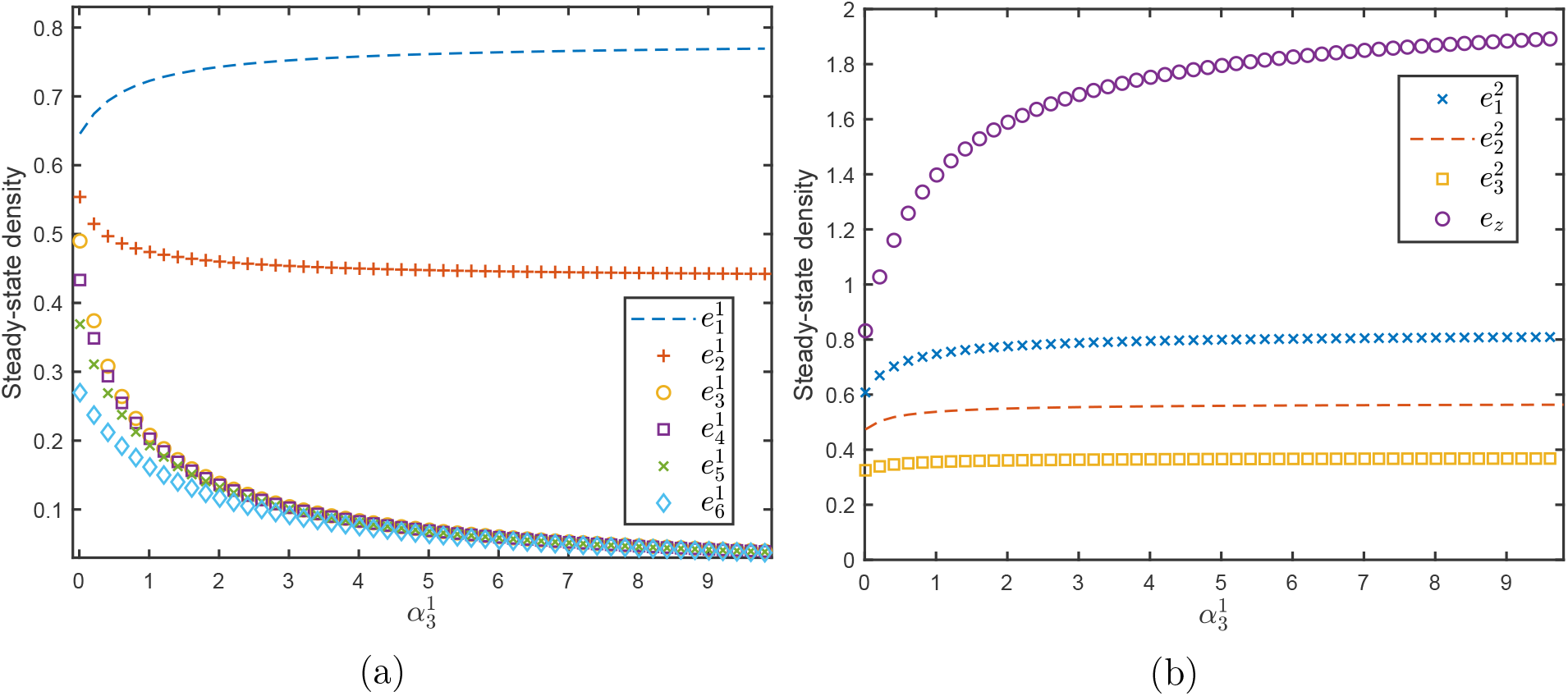
Behaviour of the RFMLKN in Example 8 as a function of 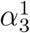 when 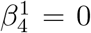: a) Steady-state values in RFMLK #1. b) Steady-state values in RFMLK #2 and the pool.

#### Example 9

Consider the RFMLKN in Example 8, but now with 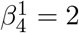. Figure 11 shows that in this case an increase in the drop-off rate 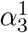 yields an increase in the steady-state values in the sites of RFMLK #1 located after the third site. This is because of an increase in the density of free ribosomes in the pool leading to more ribosomes attaching to the first RFMLK.

**Figure 11:**
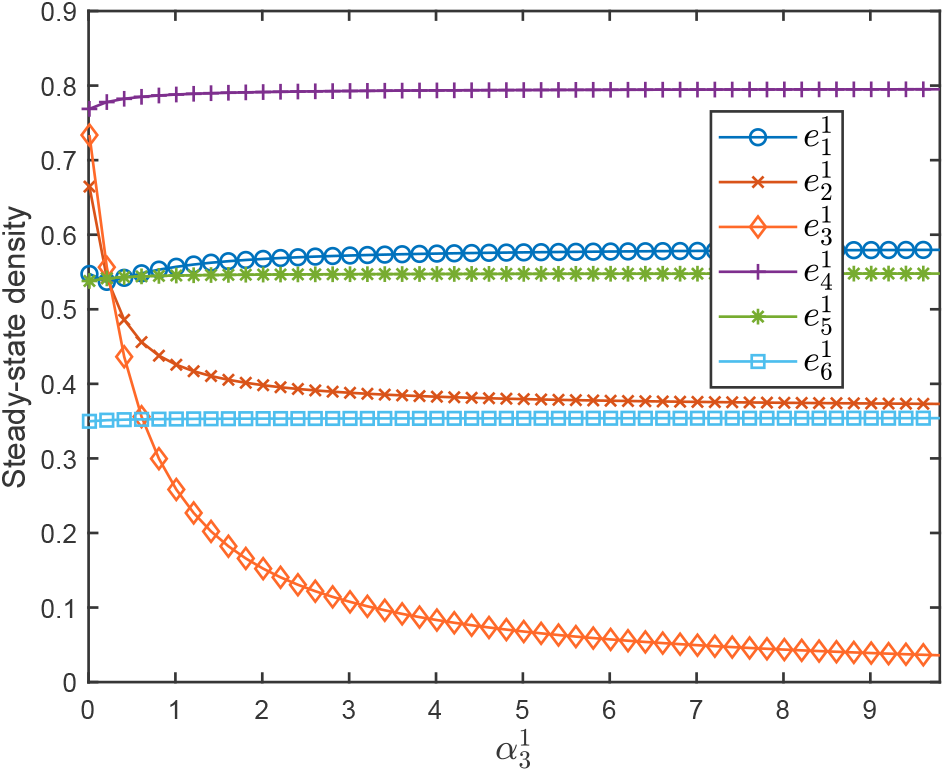
Steady-state values in RFMLK #1 in the RFMLKN in Example 9 as a function of the drop-off rate 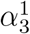.

The next result analyzes the “dual” case i.e. the affect of modifying one of the attachment rates in an RFMLK in the network.

#### Theorem 4.3

*Consider an RFMLKN with m RFMLKs of dimensions n*_*i*_, *i* = 1, …, *m, and parameters* 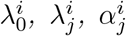, *and* 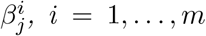, *i* = 1, …, *m and j* = 1, 2, …, *n*_*i*_. *Fix s* > 0, *and let* 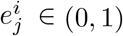, *and e*_*z*_ ∈ (0, ∞) *denote the unique steady-state density in the level set L*_*s*_ *of H. Fix k* ∈ {1, …, *n*_1_}, *and suppose that we modify the RFMLKN by changing* 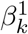 *to* 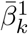, *with*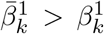. *Let* 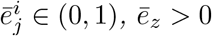, *denote the steady-state density in the modified RFMLKN. Then*

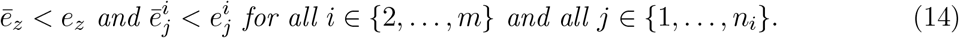

In other words, an increase in the attachment rate in one of the RFMLKs decreases the steady-state pool density and consequently decreases the density in each site in all the *other* RFMLKs.

#### Example 10

Consider an RFMLKN with *m* = 2 RFMLKs of dimensions *n*_1_ = 9 and *n*_2_ = 3, and parameters 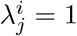, for all *i, j*, 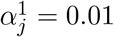, for all *j* except for 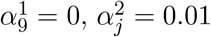, for *j* = 1, 2, 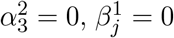 for all *j* except for 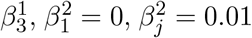 for *j* = 2, 3, and with *G*_*i*_(*z*) = *z, i* = 1, 2. The initial condition is 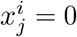, for all *i, j*, and *z*(0) = 5. We simulated this RFMLKN for several values of the attachment rate 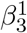. Figure 12b shows that as 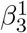 increases there is a decrease in the steady-state pool density (as more ribosomes bind to the third site of RFMLK #1) and consequently a decrease in the steady-state values in all the sites in RFMLK #2. As shown in Figure 12a, the effect of increasing 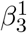 on RFMLK #1 is non-trivial. The decrease in the pool density, decreases 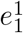. However, the increase in the attachment rate in the third site leads to more ribosomes attaching to this site and consequently a higher density in sites 3, …, 9. Also, this creates a “traffic jam” along these sites and thus increases the density in site 2 as well.

**Figure 12:**
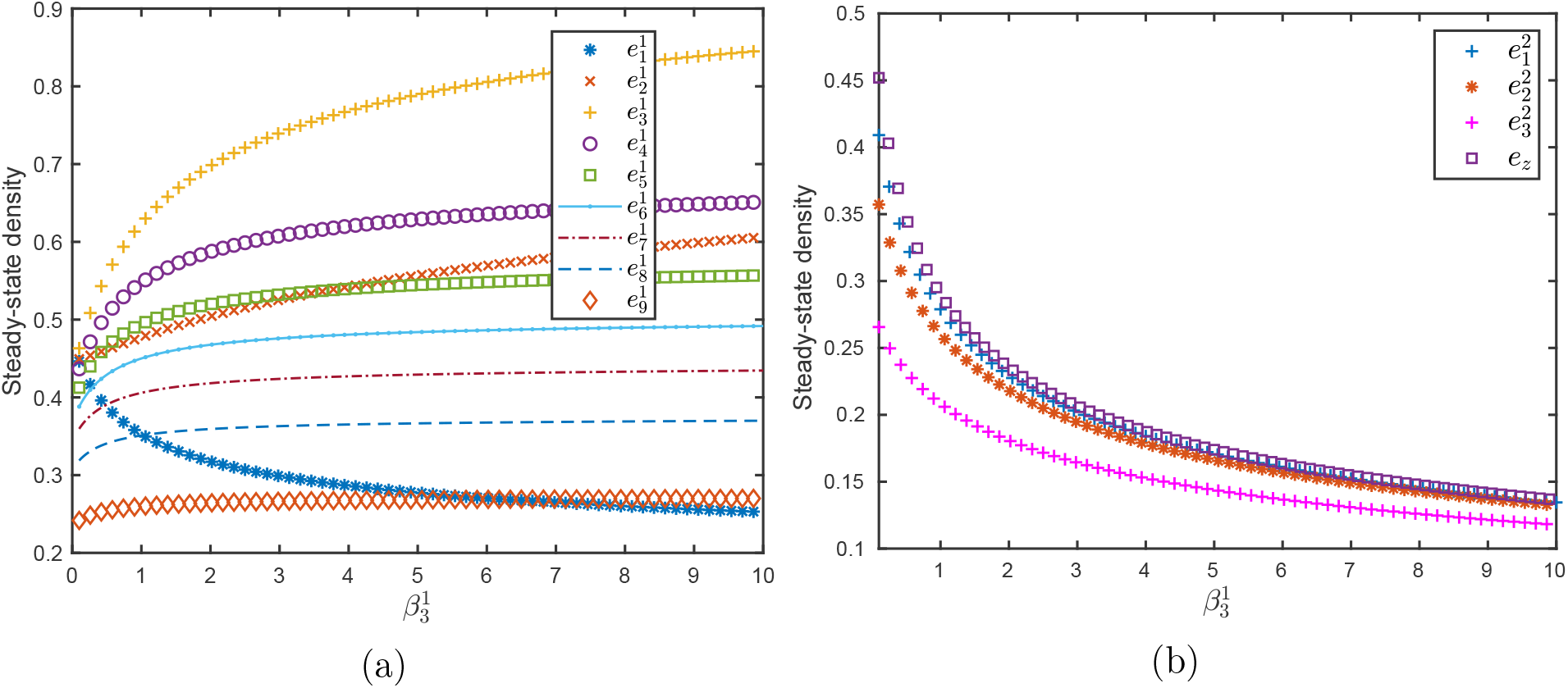
Behaviour of the RFMLKN in Example 10 as a function of the attachment rate 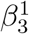: a) Steady-state densities RFMLK #1. b) Steady-state densities in RFMLK #2 and the pool.

Our last result in this subsection analyzes the effect of modifying a hopping rate in one of the RFMLKs in the network.

#### Theorem 4.4

*Consider an RFMLKN with m RFMLKs of dimensions n*_*i*_, *i* = 1, …, *m, and parameters* 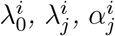 *and* 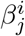, *i* = 1, …, *m, j* = 1, 2, …, *n*_*i*_. *Fix s* > 0, *and let* 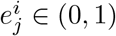, *and e*_*z*_ ∈ (0, ∞) *denote the unique steady-state density in the level set L*_*s*_ *of H. Fix k* ∈ {0, …, *n*_1_}. *Suppose that we modify the RFMLKN by changing* 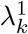 *to* 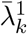, *with* 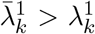. *Let* 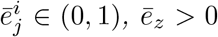 *denote the steady-state density in the modified RFMLKN. Then one of the following three cases holds. Either*

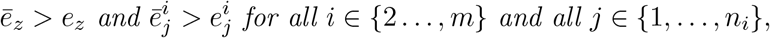

*or*

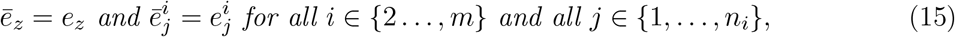

*or*

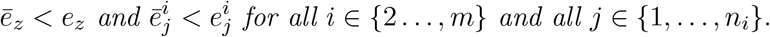

Clearly, this covers all the possible cases for the change in the pool density, and each case shows that the qualitative behaviour of all the other RFMLKs is the same.

Theorem 4.4 does not provide any information on the modified densities in the sites along RFMLK #1, as any of these densities may increase or decrease depending upon the parameters in the entire network. The next examples demonstrate this

#### Example 11

Consider an RFMLKN with *m* = 2 RFMLKs of dimensions *n*_1_ = 9 and *n*_2_ = 3, parameters 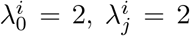, for all *i, j* except for 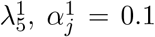, for *j* ∈ {1, 2, …, 8}, 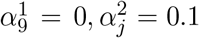, for *j* = 1, 2, 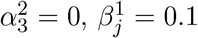 for all *j* except for 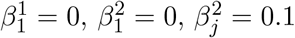 for *j* = 2, 3 and *G*_*i*_(*z*) = *z, i* = 1, 2. The initial condition is 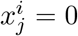, for all *i, j*, and *z*(0) = 3. We simulated this RFMLKN for various values of the elongation rate 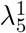. Note that when 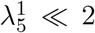 it is a bottleneck rate in RFMLK #1. Figure 13b shows that increasing 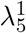 increases the pool density and thus the densities in all the sites along RFMLK #2. Figure 13a shows that increasing 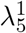 yields a decrease in sites 3, 4, 5 in RFMLK #1, but an increase in all the other sites.

**Figure 13:**
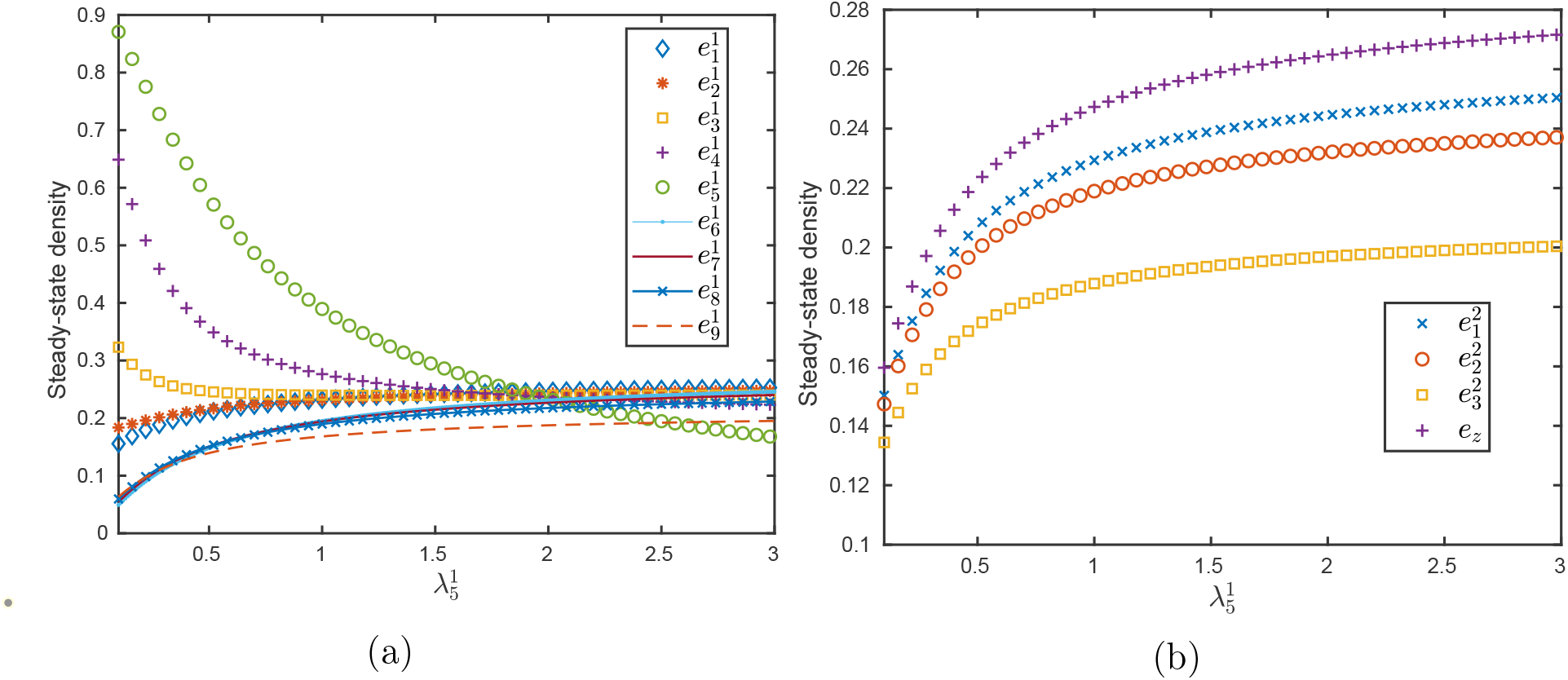
Behaviour of the RFMLKN in Example 11 as a function of the elongation rate 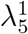: a) Steady-state densities in RFMLK #1. b) Steady-state densities in RFMLK #2 and the pool.

#### Example 12

Consider again the RFMLKN in Example 11, but now the initial condition is 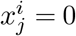, for all *i, j*, and *z*(0) = 10. In this case, Figure 14b shows that increasing the elongation rate 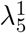 leads to a decrease in the steady-state pool density resulting in decreased steady-state densities in RFMLK #2. Figure 14a shows that in this case the effect of increasing 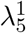 on RFMLK #1 is more intuitive: it decreases the densities in sites 1, …, 5 and increases the densities in sites 6, …, 9.

**Figure 14:**
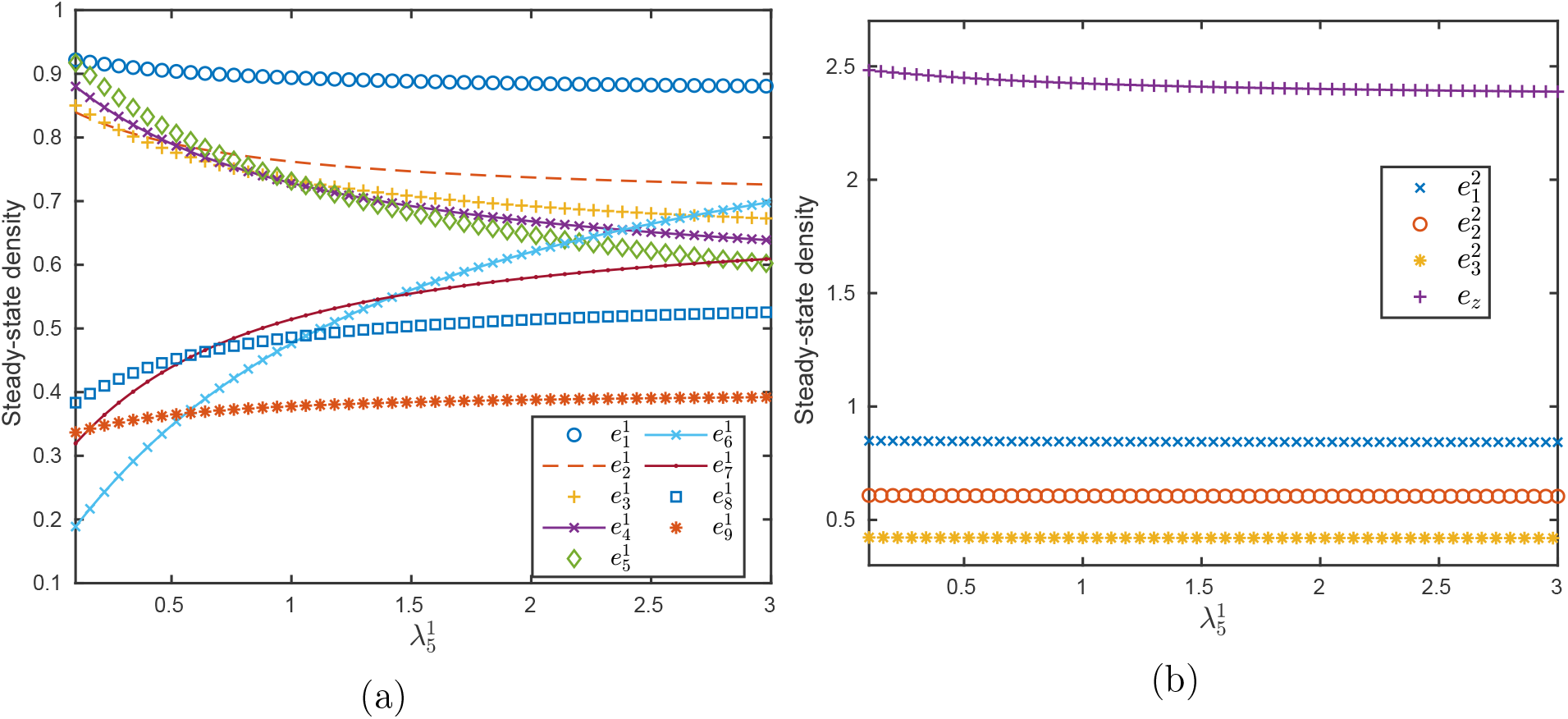
Behaviour of the RFMLKN in Example 12 as a function of the elongation rate 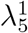: a) Steady-state densities in RFMLK #1. b) Steady-state densities in RFMLK #2 and the pool.

We now analyze several additional mathematical properties of the RFMLKN.

### 4.8 Strong monotonicity

Recall that the dynamical system 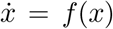 is called cooperative if for any two initial conditions *a, b* with *a* ≤ *b* we have *x*(*t, a*) ≤ *x*(*t, b*) for all *t* ≥ 0 [54]. In other words, the flow preserves the (partial) ordering between the initial conditions. The next result shows that the RFMLKN is cooperative.

#### Proposition 4.8

*Consider the RFMLKN. Fix two initial conditions a, b* ∈ Ω *with a* ≤ *b. Then*

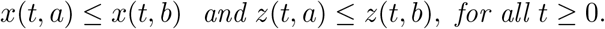

*If, furthermore, a* ≠ *b then*

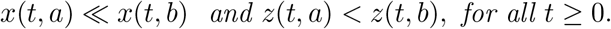

In other words, if we consider two initial conditions where the first corresponds to a smaller density in each site in each RFMLK and in the pool, then the corresponding solutions will satisfy the same relation at any time *t* ≥ 0.

The next subsection shows that the flow of the RFMLKN is a non-expansive mapping. For a vector *v* ∈ ℝ^*n*^, let 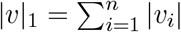 denote the *L*_1_ norm of *v*.

### 4.9 Non-expansion

In a contractive system, all solutions converge exponentially to one another. Since the RFMLKN admits more than one equilibrium, it is not a contractive system with respect to any norm. However, the next result shows that the *L*_1_ distance between any two trajectories is non-expansive, i.e. it is bounded by the distance *L*_1_ between the initial conditions.

#### Proposition 4.9

*Consider the RFMLKN. Fix a, b* ∈ Ω. *Then*

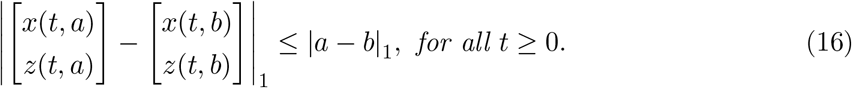

In particular, the difference between two “close” ribosomal density profiles will remain close for all *t* ≥ 0.

Fix *a* ∈ Ω and let 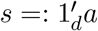. Setting 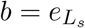 in (16) yields

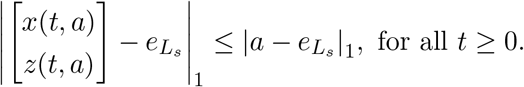

In other words, the convergence to the equilibrium 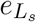 is monotone in the sense that the *L*_1_ distance can only decrease with time.

### 4.10 Entrainment

Many biological processes are excited by a periodic input. Proper functioning often requires entraining to the excitation, that is, converging to a periodic pattern with the same period as the excitation. A typical example is the ability of cells to coordinate their growth with the periodic cell-cycle division process. Translation seems to play an important role in this process. It is known for example that expression of the human translation initiation factor eIF3f peaks twice in the cell cycle: in the S and the M phases [64].

Entrainment is also important in the context of synthetic biology, for example, in designing a biological network that is coordinated by a single oscillator producing a common “clock signal” [68].

To study entrainment in the RFMLKN, assume that the parameters 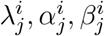 in all the RFMLKs are not constant, but are time-varying functions, that are all jointly periodic with a period *T >* 0. More precisely, we assume that

- There exists a (minimal) *T >* 0 such that all the non-negative time-varying rate functions 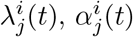 and 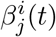 are T-periodic.
- There exists 0 *< δ*_1_ ≤ *δ*_2_ such that 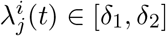, for all *i, j* and all *t* ∈ [0, *T*).

We then refer to the network as the periodic RFM with Langmuir kinetics network (PRFMLKN). Note that a constant function is *T* -periodic for any *T*, so for example if one parameter in the network is *T* -periodic and all the others are constant then our assumptions hold.

The next result shows that the PRFMLKN entrains.

#### Theorem 4.5

*Consider the PRFMLKN. Fix s* ≥ 0. *There exists a unique function ϕ*_*s*_ : ℝ_+_ → int(Ω), *that is T -periodic, and for any initial condition* 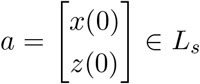, *the solution* 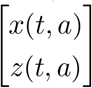 *of the PRFMLKN converges to ϕ*_*s*_ *as t* → ∞.

In other words, if the rates are *T* -periodic then all the densities in the mRNAs and the pool converge to a periodic pattern with period *T*, and thus so will the protein production rate in every mRNA. In particular, if a single parameter in one of the RFMLKs is *T* -periodic and all the other parameters are constant then the network entrains. Roughly speaking, this can be explained as follows. The *T* -periodic parameter will generate *T* -periodic variations in the pool density and this will generate *T* -periodic patterns in all the mRNA densities, as the pool feeds all the mRNAs. Again, this demonstrates the intricate coupling generated by the competition for shared resources.

#### Example 13

Consider a network with *m* = 2 RFMLKs, of dimensions *n*_1_ = 2 and *n*_2_ = 3, and with *G*_*i*_(*z*) = tanh(*z*), *i* = 1, 2. All the rates in the network are equal to one, except for 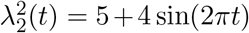. Thus all the rates in the network are periodic with a common minimal period *T* = 1. The initial condition is 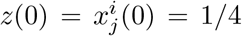 for all *i, j*. Figure 15 depicts the state-variables and the pool density as a function of *t*. Note that all the densities converge to a periodic pattern with period one. Note also that since the total number of ribosomes in conserved, maximal peaks in the density along the RFMLKs corresponds to minimal peaks in the pool density, and vice-versa.

**Figure 15:**
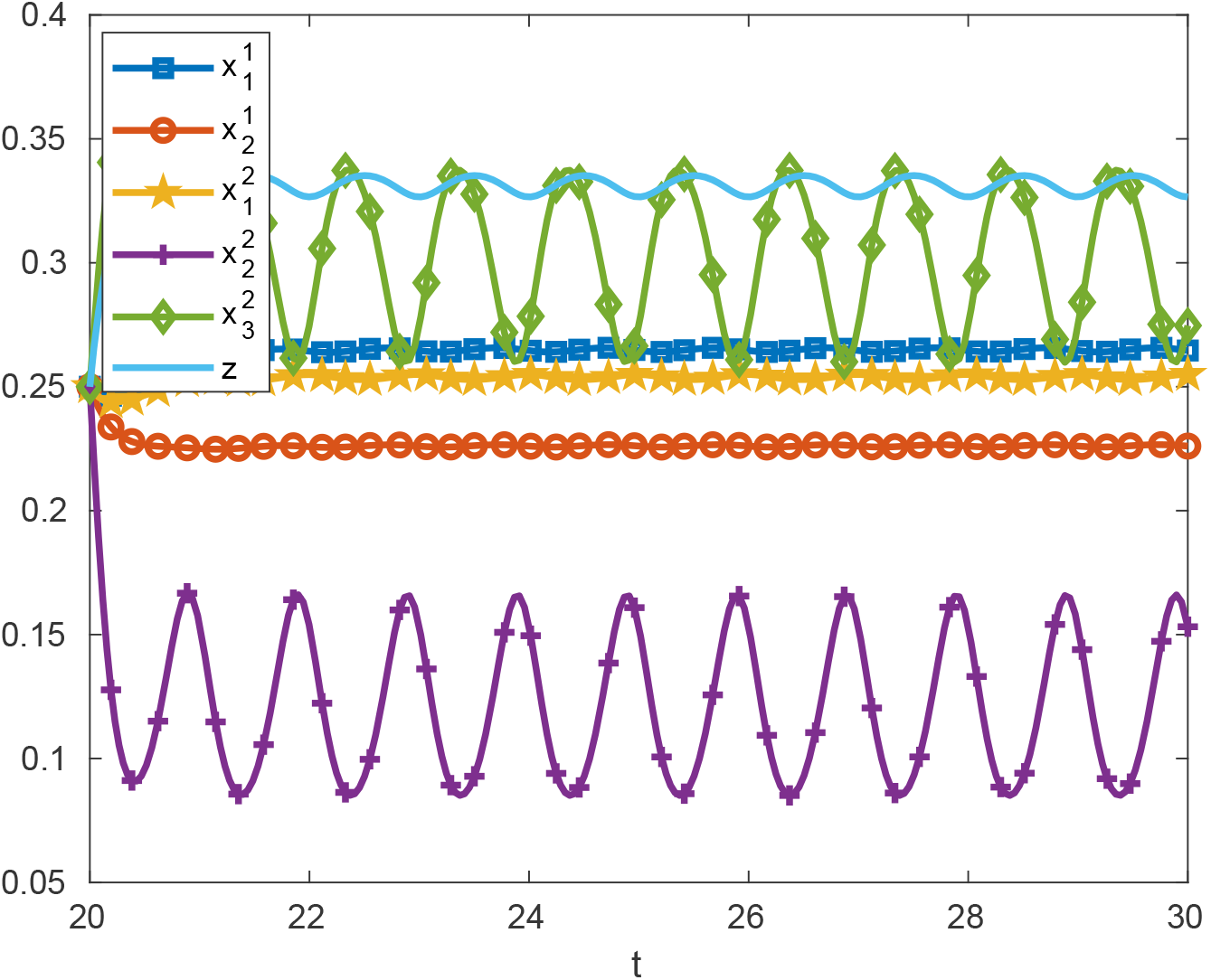
Trajectories of PRFMLKN in Example 13 as a function of time.

## 5 Discussion

We derived and analyzed a novel and general network model of ribosome flow during large-scale translation in the cell. This model encapsulates important cellular properties like ribosome drop-off, ribosome attachment at IRESs, and competition for a finite pool of free ribosomes. We analyzed the model using tools from systems and control theory, including contraction theory, and the theory of cooperative dynamical systems.

The new model is an irreducible cooperative dynamical system admitting a first integral (the total density of ribosomes in the network). This implies that the system admits a continuum of linearly ordered equilibrium points, and that every trajectory converges to an equilibrium point. The system is also on the “verge of contraction” with respect to the *L*_1_ norm. In addition, we proved that if one or more of the rates in the network are time-varying periodic functions with a common period *T*, then the densities along all the mRNAs and in the pool converge to a periodic solution with period *T*, i.e. the system entrains to a periodic excitation.

An important question is the sensitivity of the network steady-state to variations in the mRNA parameters and the density of free ribosomes. We thoroughly analyzed this problem, and showed that a modification of a bio-physical property in a specific mRNA has two implications. First, via competition it affects translation in all the other mRNAs in an intuitive manner: if the pool steady-state density increases [decreases] then the density in all other sites in all other mRNAs increases [decreases]. Second, and perhaps surprisingly, it is almost impossible to predict what will be the effect on the densities and protein production rate in the mRNA that is modified, as this depends in a non-trivial way on the interactions between all the mRNAs and the pool. For example, an increase in the drop-off rate in a specific site in an mRNA may increase the pool density, thus increasing the attachment rates along this mRNA and leading to an increase in the density in some sites along this mRNA.

These results highlight that analyzing the effect of any bio-physical property on translation in the cell must take into account the intricate effects of competition, especially when the competition for shared resources plays a major role, e.g. under stress conditions.

We believe that the new model presented here provides a powerful framework for analyzing and re-engineering the translation process. One possible avenue for further research is in developing a quantitative and qualitative understanding of how viral mRNAs hijack the translation machinery and, in particular, whether the indirect effects of competition are enough to hamper the host’s immune response.

## 6 Appendix: Proofs

### 6.1 Proofs of results for the RFMLK

We begin by writing the RFMLK with input and output in the form:

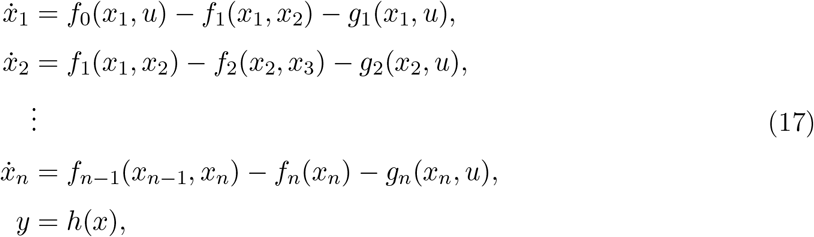

where *u* : ℝ _+_ → ℝ _+_ is a scalar input function that takes non-negative values for any time *t* ≥ 0, *y* : ℝ _+_ → ℝ _+_ is a scalar non-negative output function, and

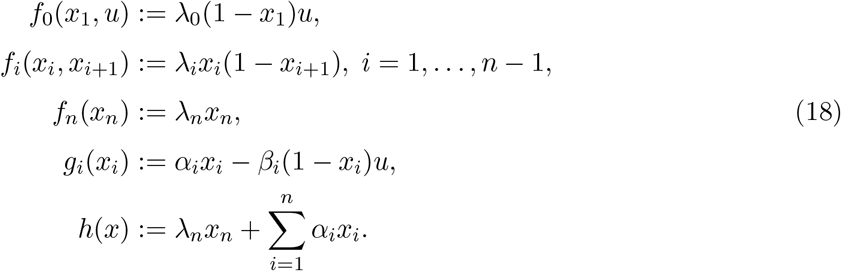

The parameters satisfy *λ*_*i*_ > 0, *α*_*i*_ ≥ 0, and *β*_*i*_ ≥ 0 for all *i*. Recall that every *x*_*i*_ takes values in the interval [0, 1], so the state-space of the RFMLK is *C*^*n*^ := [0, 1]^*n*^.

#### 6.1.1 Proof of Prop. 4.1

Fix *δ* > 0. We will show that for any sufficiently small Δ > 0 there exists *K* = *K*(*δ*, Δ) > 0 such that for every *k* ∈ {1, …, *n*} and every *t* ≥ 0 the condition

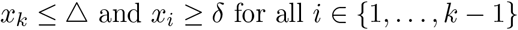

implies that

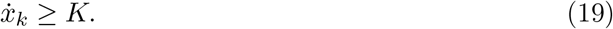

For *k* = 1 the condition is simply *x*_1_ ≤ Δ, and then

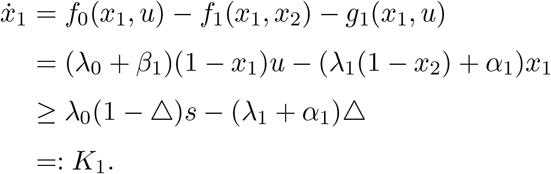

Note that *K*_1_ ≥ *λ*_0_*s/*2 > 0 for any Δ > 0 sufficiently small. For *k* ∈ {2, …, *n* − 1} we have

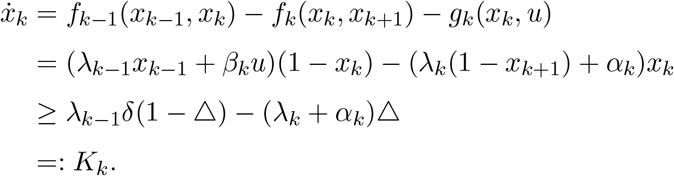

Note that *K*_*k*_ ≥ *λ*_*k*−1_*δ/*2 > 0 for any Δ > 0 sufficiently small. For *k* = *n* we have

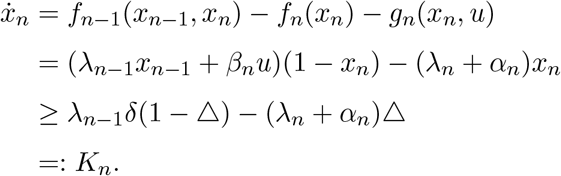

Note that *K*_*n*_ ≥ *λ*_*n*−1_*δ/*2 > 0 for any Δ > 0 sufficiently small.

We conclude that (19) holds for *K* := min{*K*_1_, …, *K*_*n*_} ≥ min{*λ*_0_, …, *λ*_*n*_} min{*s, δ*}*/*2 > 0. By [36, Lemma 1], this implies that for any *τ >* 0 there exists *ϵ* _1_ = *ϵ* _1_(*τ*) > 0, with *ϵ* _1_(*τ*) → 0 as *τ* → 0, such that for any *a* ∈ *C*^*n*^ the solution of (17) satisfies

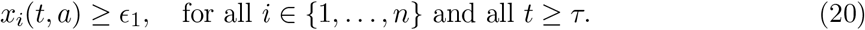

Let *z*_*i*_ := 1 − *x*_*n*+1−*i*_, *i* = 1, …, *n*. Then (17) gives

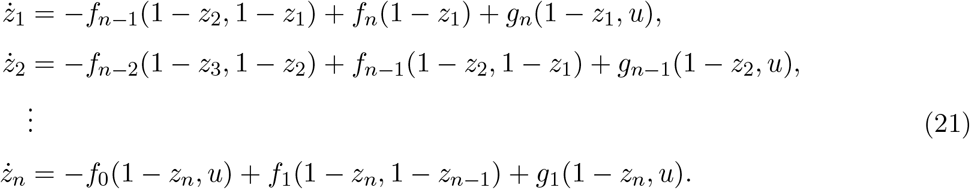

It is not difficult to verify that this system also satisfies condition (19), so by [36, Lemma 1], for any *τ >* 0 there exists *ϵ* _2_ = *ϵ* _2_(*τ*) > 0, with *ϵ* _2_(*τ*) → 0 as *τ* → 0, such that for any *a* ∈ *C*^*n*^ the solution of (21) satisfies

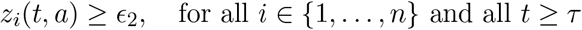

Combining this with (20) completes the proof of Prop. 4.1.

#### 6.1.2 Proof of Prop. 4.2

Let *f* denote the vector field in (17), and let 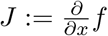 denote its Jacobian. Then

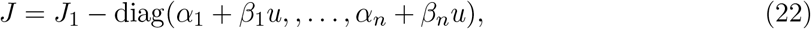

where

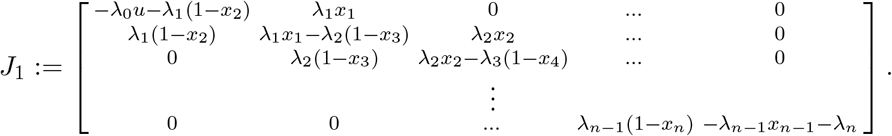

Note that *J*_1_ is the Jacobian of an RFM with a time-varying initiation rate *λ*_0_*u*(*t*) ≥ *λ*_0_*s*, and that *α*_*i*_ + *β*_*i*_*u*(*t*) ≥ 0 for all *t*. Note also that for any *x* ∈ int(*C*^*n*^), all the entries in the super- and sub-diagonal of *J*_1_ are positive, so in particular *J*_1_ (and thus *J*) is irreducible.

Fix *a, b* ∈ *C*^*n*^ and *τ >* 0. By Proposition 4.1, there exists *ϵ* = *ϵ* (*τ*) > 0, such that

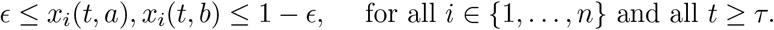

Now arguing as in the proof of [36, Theorem 4] completes the proof of Prop. 4.2.

#### 6.1.3 Proof of Prop. 4.3

The RFMLK with a constant input *u*(*t*) ≡ *s >* 0 is a time-invariant system that maps the convex and compact state-space [0, 1]^*n*^ to itself. Hence, it admits an equilibrium *e*^*s*^. Prop. 4.1 implies that *e*^*s*^ ∈ (0, 1)^*n*^. Prop. 4.2 implies that any solution converges to *e*^*s*^, and this completes the proof of Prop. 4.3.

#### 6.1.4 Proof of Prop. 4.4

It follows from (22) that *J* is a Metzler matrix, i.e. every off-diagonal entry of *J* is non-negative. Also,

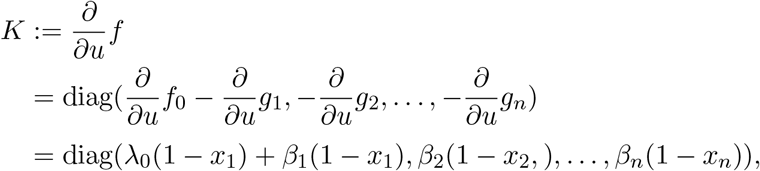

so every entry of *K* is non-negative. The results in [31] imply that the RFMLK is a monotone control system, so (7) holds. Now the definition of the output *y* implies (8), and this completes the proof of Prop. 4.4.

#### 6.1.5 Proof of Prop. 4.5

We already know that the limits 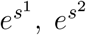, and *e* := lim_*t*→∞_ *x*(*t, a, v*) exist. By monotonicity, *x*(*t, a, u*) ≤ *x*(*t, a, v*), for all *t* ≥ 0, and taking the limit as *t* → ∞ gives 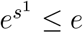. Since the system is contractive, 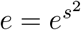, and this completes the proof of Prop. 4.5.

### 6.2 Proofs of results for the RFMLKN

For the sake of simplicity and to avoid cumbersome notation, we provide proofs of the theoretical results when *m* = 2, i.e. a network with two RFMLKs connected via a pool of free ribosomes (the proofs when *m >* 2 are very similar). We write the first RFMLK as

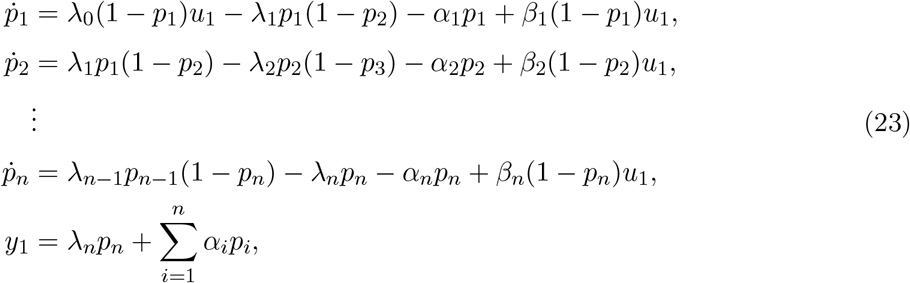

and the second as

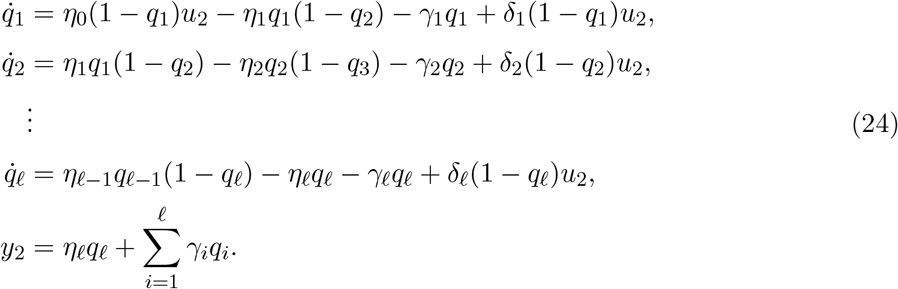

The inputs to the RFMLKs are functions of the pool density

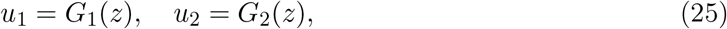

and the pool dynamics is

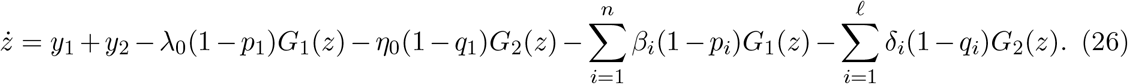

The state-space of this network is

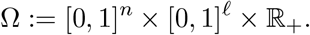

Note that 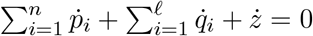, so

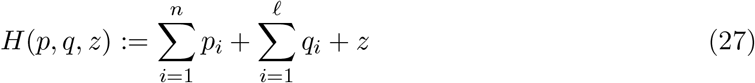

is a first integral of the dynamics, that is, *H*(*p*(*t*), *q*(*t*), *z*(*t*)) ≡ *H*(*p*(0), *q*(0), *z*(0)). In other words, the total density of ribosomes in the network is conserved. For any *s* ≥ 0, we define the *s* level set of the first integral *H* by

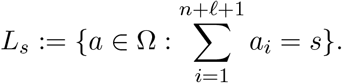

Thus, *L*_*s*_ includes all the states in Ω with total density of ribosomes equal to *s*. Note that for *s* = 0, *L*_0_ = {0} and the dynamics emanating from zero remains in zero for all time *t* ≥ 0. therefore, we will always consider *L*_*s*_ with *s >* 0.

#### 6.2.1 Proof of Prop. 4.7

We now restate and prove the persistence result in Prop. 4.7 for the case *m* = 2.

##### Proposition 6.1

*Consider the RFMLKN with m* = 2. *Fix s >* 0. *Then for any τ >* 0 *there exists ϵ* (*τ*) > 0, *with ϵ* (*τ*) → 0 *as τ* → 0, *such that for any initial condition in L*_*s*_ *and any t* ≥ *τ the solution of the RFMLKN satisfies*

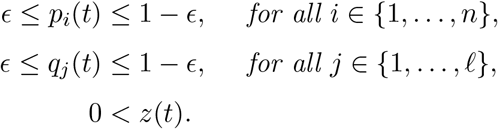

**Proof:** It is useful to denote *p*_0_ := *z, p*_*n*+1_ = 0, and *p*_−1_ := *p*_*n*_. Using the fact that *y*_2_ ≥ 0 yields

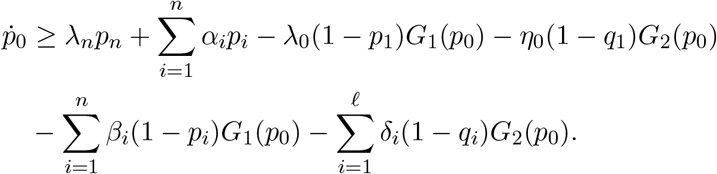

We now show that the system with state-variables *p*_0_, …, *p*_*n*_ satisfies t he c yclic boundary-repelling (CBR) condition in [33, Lemma 1], that is, for any *δ >* 0 and any sufficiently small Δ > 0, there exists *K* = *K*(*δ*, Δ) > 0 such that for each *k* = 0, …, *n* and each *t* ≥ 0 the condition

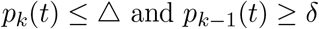

implies that

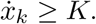

Indeed, for *k* = 0 we have

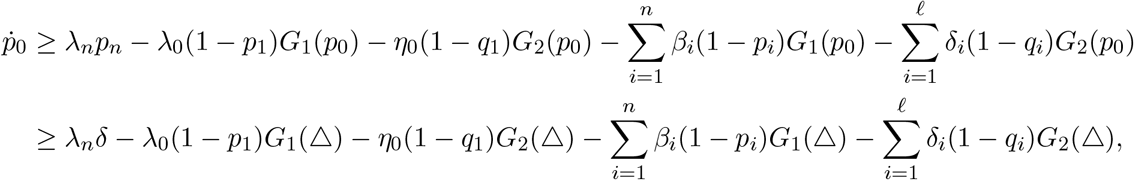

and since *G*_*i*_(0) = 0 and *G*_*i*_ is continuous, 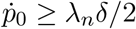 for all Δ > 0 sufficiently small.

For *k* ∈ {1, …, *n*} we have

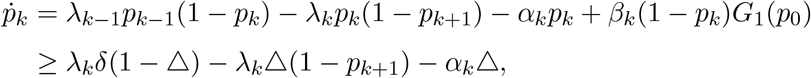

so 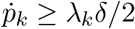 for all Δ > 0 sufficiently small.

It is also straightforward to verify that if *p*_*k*_(*t*) > 0 for some *k* ∈ {0, …, *n*} and *t >* 0 then *p*_*k*_(*r*) > 0 for all *r* ≥ *t*. It now follows from [33, Lemma 1] that for any *τ >* 0 there exists *ϵ* (*τ*) > 0, with *ϵ* (*τ*) → 0 as *τ* → 0, such that for any initial condition [*p*_0_(0) … *p*_*n*_(0)]^*T*^= ≠ 0 and any *t* ≥ *τ* the solution of the RFMLKN satisfies

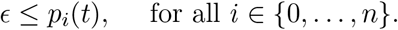

Using a similar argument for the *q* system proves that for any *t* ≥ *τ*,

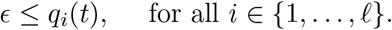

Finally, arguing as in the proof of Prop. 4.1 completes the proof of Prop. 6.1. □

Our next goal is to prove the sensitivity results for the RFMLKN. It is useful to first write equations describing the steady-state of the RFMLK for a constant input *u*(*t*) ≡ *v*, with *v >* 0. By (3),

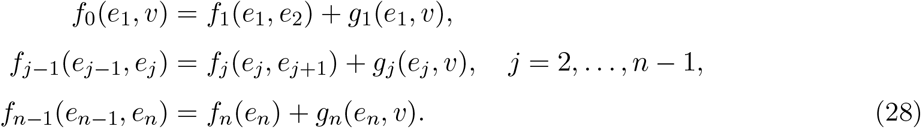

This yields

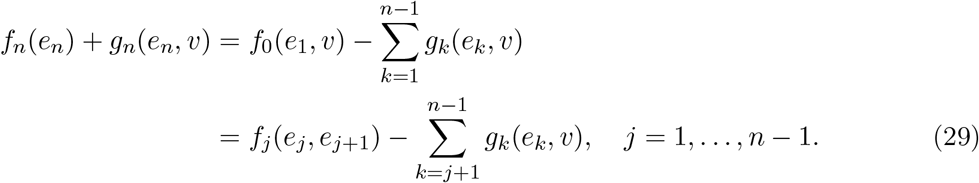

Substituting the expressions for the *f*_*i*_s and *g*_*i*_s yields the following result.

##### Proposition 6.2

*Consider the steady-state of the RFMLK with u*(*t*) ≡ *v, where v >* 0. *Then for any k* = 1, …, *n* − 1, *we have*

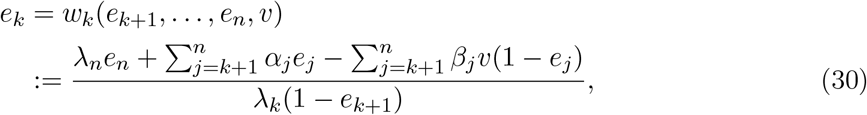

*and*

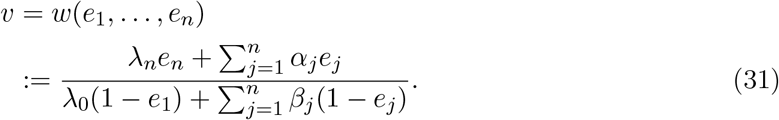

Note that the function *w*_*k*_ is increasing in *e*_*k*+1_, …, *e*_*n*_ (and strictly increasing in *e*_*k*+1_, *e*_*n*_), and is decreasing in *v*. Also, the function *w* is increasing in *e*_1_, …, *e*_*n*_ (and strictly increasing in *e*_1_, *e*_*n*_).

#### 6.2.2 Proof of Theorem 4.1

It is clear that for *s* = 0, *L*_0_ = {*e*^0^}, with *e*^0^ = 0, and all trajectories converge to *e*^0^. Fix *s >* 0. The Jacobian of the RFMLKN with *m* = 2 is

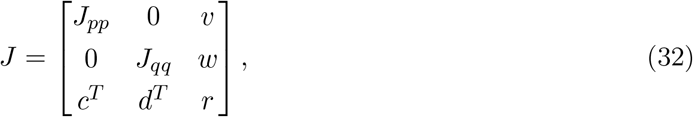

where *J*_*pp*_ is the Jacobian of an RFMLK with state-variables *p*_1_, …, *p*_*n*_, rates *λ*_*i*_, *α*_*i*_, *β*_*i*_ and input *u* = *G*_1_(*z*) (see (22)), *J*_*qq*_ is the Jacobian of an RFMLK with state-variables *q*_1_, …, *q* _*ℓ*_, rates *η*_*i*_, *γ*_*i*_, *δ*_*i*_ and input *u* = *G*_2_(*z*) (see (22)),

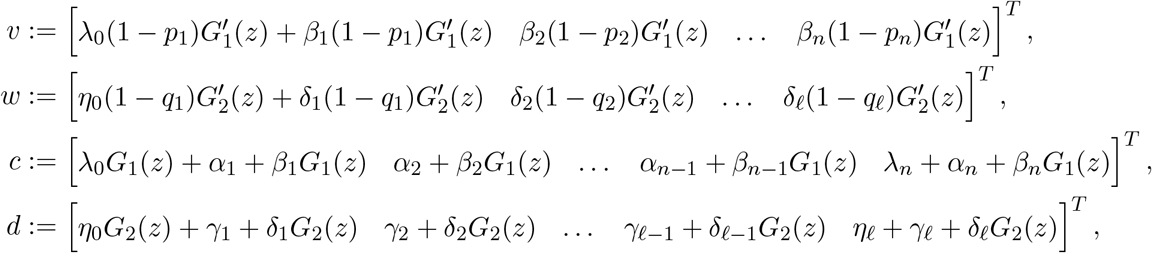

and

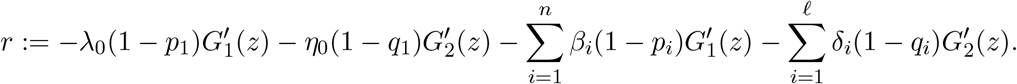

These equations imply that for any [*p q z*] ^*T*^ ∈ Ω the Jacobian matrix *J* is Metzler, so the RFMLKN is a cooperative dynamical system. Furthermore, for any [*p q z*]^*T*^ ∈ int(Ω) all the entries on the super- and sub-diagonals of *J*_*pp*_, *J*_*qq*_ are positive, and so are the first entry in *v, w*, and the first and last entry in *c, d*. This implies that for any [*p q z*] ^*T*^ ∈ int(Ω) the matrix *J* is irreducible. Combining this with Prop. 6.1 and the results in [56] on strongly cooperative dynamical systems with a first integral completes the proof of Theorem 4.1.

#### 6.2.3 Proof of Theorem 4.2

We again prove for the special case *m* = 2, i.e. a network with two RFMLKs and a pool. Let 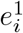, *i* ∈ {1, …, *n*}, denote the steady-state in the first RFMLK, and 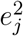, *j* ∈ {1, …, *ℓ*}, denote the steady-state in the second RFMLK.

Since the initial condition remains the same, we have:

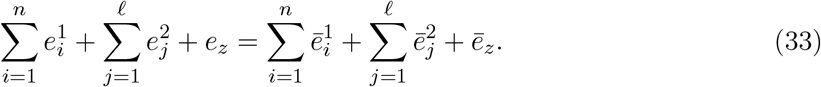

We prove that *e*_*z*_ *< ē*_*z*_ by contradiction. We consider two cases: *e*_*z*_ = *ē*_*z*_; and *e*_*z*_ *> ē*_*z*_, and show that each of these cases yields a contradiction.

**Case 1**. Assume that *e*_*z*_ = *ē*_*z*_. In this case, there is no change in the input and parameter values in the second RFMLK, so 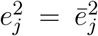 for all *j* = 1, …, *ℓ*. Consider the first RFMLK. Since the steady-state input to this RFMLK remains the same, Prop. 7 in Ref. [35], that states that increasing any of the detachment rates (without changing any other parameter) in the RFMLK decreases all the steady-state densities, implies that 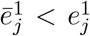 for all *j*. However, this contradicts (33).

**Case 2**. Assume that

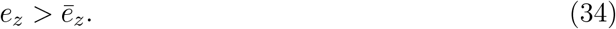

In other words, after increasing 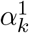 to 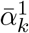, the steady-state input to each RFMLK is decreased. Then 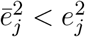 for all *j* = 1, …, *ℓ*. Combining this with (33) implies that 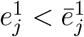 for at least one index *j*. Let *s* ∈ {1, …, *n*} be the *maximal index* such that

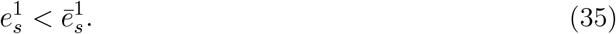

Applying (30) inductively implies that 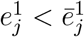 for any *j* ∈ {*s, s* − 1, …, 1} (note that it follows from (30) that increasing 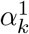 can only further increase the corresponding steady-state value 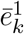). Suppose that

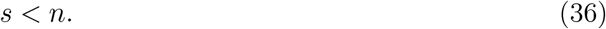

Then

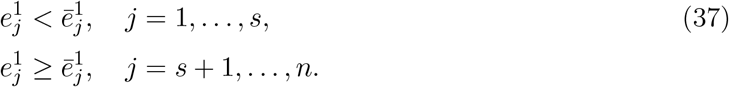

By (30),

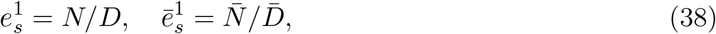

where

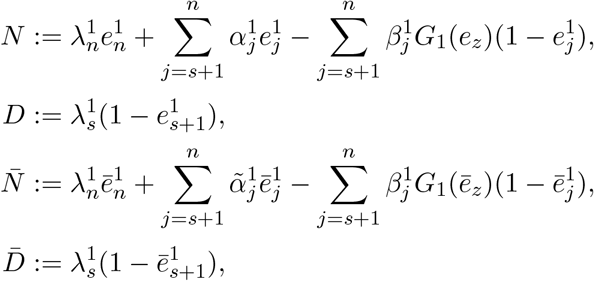

where 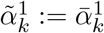, and 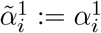, for all *i* ≠ *k*. Note that (37) implies that

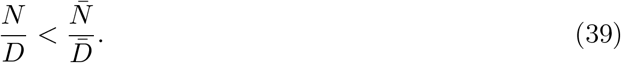

By the first equation in (29),

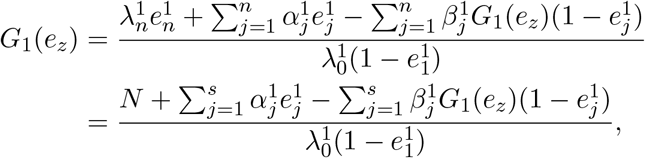

and thus

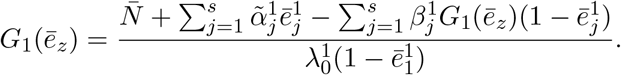

Combining this with (34) and (37) implies that if 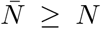 then *G*_1_(*ē*_*z*_) ≥ *G*_1_(*e*_*z*_), but this contradicts (34), so we conclude that 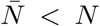. Now (39) implies that 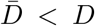, i.e. 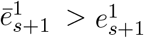. However, this contradicts (37), so we conclude that (36) cannot hold, that is, *s* = *n*. Applying (30) inductively implies that

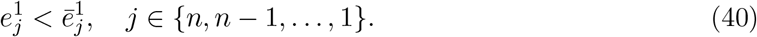

Eq. (31) gives

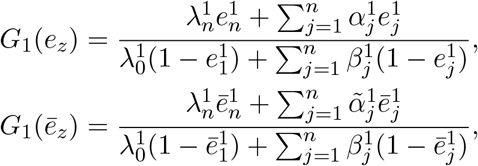

so *G*_1_(*e*_*z*_) *< G*_1_(*ē*_*z*_). This contradicts (34), and we conclude that Case 2 is impossible. This completes the proof of Theorem 4.2.

#### 6.2.4 Proof of Theorem 4.3

The proof is similar to the proof of Theorem 4.2 above and is thus omitted.

#### 6.2.5 Proof of Theorem 4.4

Clearly, we have three possible cases: *ē*_*z*_ = *e*_*z*_, or *ē*_*z*_ *> e*_*z*_, or *ē*_*z*_ *< e*_*z*_. In the first case, the input to each RFMLK satisfies *G*_*i*_(*ē*_*z*_) = *G*_*i*_(*e*_*z*_). Since the parameters in all the RFMLKs, except for RFMLK #1, are unchanged, we conclude that (15) holds. The analysis in the second and third cases is similar. This completes the proof of Theorem 4.4.

#### 6.2.6 Proof of Prop. 4.8

It was shown in the proof of Theorem 4.1 that the RFMLKN is a strongly cooperative dynamical system and the results in Prop. 4.8 follow immediately.

#### 6.2.7 Proof of Prop. 4.9

Recall that the Jacobian *J* of the RFMLKN (with *m* = 2) is given in (32). This matrix is Metzler, and a calculation shows that the sum of every column of *J* is zero. Hence, the *L*_1_ matrix measure of *J* is zero, and this implies (16).

#### 6.2.8 Proof of Theorem 4.5

Write the PRFMLKN as 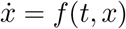. Then *f* (*t, z*) = *f* (*t*+*T, z*) for all *t* ≥ 0 and *z* ∈ Ω, and *H*(*x*) is a first integral of the dynamics. Now Theorem 4.5 now follows from the results in [55]. The fact that *ϕ*_*s*_ ∈ int(Ω) follows from the persistence result in Prop. 4.7.

